# FABP5 in Skin Macrophages Mediates Saturated Fat-Induced IL-1β Signaling in Obesity-Associated Psoriasis Development

**DOI:** 10.1101/2024.12.16.628705

**Authors:** Jianyu Yu, Jiaqing Hao, Matthew S. Yorek, Xingshan Jiang, Anthony Avellino, Shanshan Liu, Xiaochun Han, Jonathan Shilyansky, Zhaohua Wang, Ali Jabbari, Bing Li

## Abstract

High fat diet (HFD)-induced obesity increases the risk and severity of psoriasis. However, the immunoregulatory effects of different HFD-induced obesity on psoriasis pathogenesis remains poorly understood. Here, mimicking human dietary fat profiles, four HFDs – saturated, monounsaturated, omega-6 and omega-3 fats – were designed and used to induce obesity in mice. Despite comparable obesity levels across groups, only the saturated HFD exacerbated imiquimod (IMQ)-induced psoriasis. This exacerbation correlated with elevated levels of IL-1β-producing macrophages, IL-17A-producing γδ T cells, and neutrophils within psoriatic lesions. Mechanistically, saturated fatty acids (FAs) promoted IL-1β/IL-17 signaling via fatty acid-binding protein 5 (FABP5)-mediated mitochondrial FA oxidation and extracellular ATP release in skin macrophages. Deletion of FABP5, either globally or specifically in macrophages, attenuated IL-1β/IL-17A signaling and alleviated IMQ-induced psoriasis. These findings identify FABP5 as a key mediator of saturated HFD-driven psoriasis via the IL-1β/IL-17 axis, offering insights into the interplay between dietary fats, obesity and psoriasis.

**Highlights:**

1. Saturated, but not unsaturated, high-fat diets (HFDs) drive the development of obesity-associated psoriasis.
2. Saturated fats enhance IL-1β/IL-17 signaling in psoriatic skin.
3. FABP5 mediates saturated fat-induced IL-1β signaling by promoting ATP production and release.
4. Deletion of FABP5 in macrophages alleviates the development of saturated HFD-associated psoriasis.

**Graphic Abstract:** 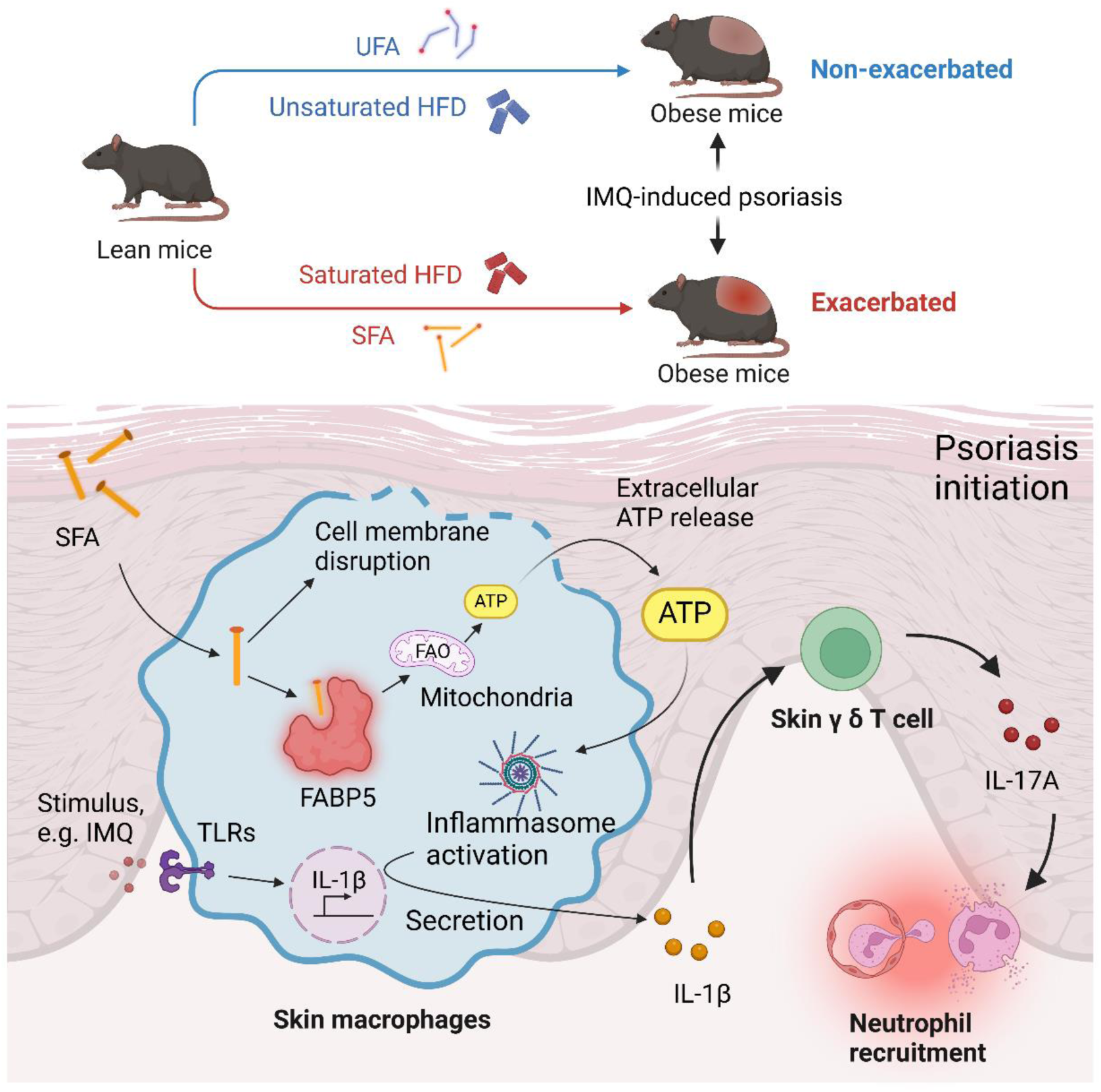

## Introduction

Fats are essential to maintain cellular function in humans^1^. Dietary fatty acids (FAs) are categorized as saturated (SFA), monounsaturated (MUFA), and polyunsaturated fatty acids (PUFA), depending on the number of double bonds in their carbon chains^2^. Despite the variety of dietary FA sources, overconsumption of high-fat diet (HFD) has been implicated in the global obesity epidemic^3^. Emerging evidence, including findings from our group, suggests that the composition of dietary fats drives distinct lipid-mediated responses in various obesity-associated diseases, including chronic inflammation, cardiovascular disease, type II diabetes, and cancer^4–7^.

Psoriasis is chronic, immune-mediated skin inflammation driven by complex interactions between immune cells and keratinocytes^8^. Clinical and animal studies have consistently shown that obesity increases both the risk and severity of psoriasis^9–11^. Diet-induced obesity (DIO) by excessive HFD intake, promotes proinflammatory immune cell infiltration and upregulates cytokines and chemokines in the skin^12,13^. However, HFDs consumed in daily life or employed in research are not homogeneous, containing distinct proportions of SFAs, MUFAs and PUFAs^14–16^. Recent studies indicate that various skin cell types exhibit differential responses to individual FAs, activating distinct lipid-mediated signaling pathways^4,17,18^. Despite these findings, the influence of specific FA compositions in HFDs on the development of obesity-related psoriasis remains poorly understood.

Macrophages, including CD207^+^ epidermal macrophages and CD207^-^ dermal macrophages, are key regulators of tissue homeostasis and immune responses, performing functions such as phagocytosis, antigen presentation, and cytokine production^19^. Emerging evidence highlights the heterogeneity of macrophages in dietary lipid uptake and metabolism^20,21^, which influences their contributions to obesity-related diseases, including cancer and chronic inflammation^4,5,22^. For instance, saturated fats drive tumor-associated macrophage activation, promoting breast cancer progression^5^, whereas unsaturated fats activate IL-36 signaling in skin macrophages, leading to alopecia in murine models^4^. Despite these advances, how skin macrophages metabolize and respond to specific dietary lipids, and how these processes influence their roles in obesity-related psoriasis, remain largely undefined.

Lipids are insoluble in aqueous environments, requiring specific mechanisms for their cellular utilization. Fatty-acid binding proteins (FABPs) have evolved to facilitate the solubilization and transport of FAs, playing a central role in regulating lipid metabolism and cellular functions^23^. The FABP family includes at least nine members, each with distinct distribution patterns. For example, FABP5, also known as epidermal FABP (E-FABP) is primarily expressed in skin tissue, while FABP4, or adipose FABP (A-FABP), is predominantly found in adipose tissue^23^. FABP5 is critical for maintaining keratinocyte differentiation^24^, and its dysregulation has been implicated in skin carcinoma and psoriasis^25^ ^26^. Previous research from our group demonstrated that keratinocyte FABP5 interacts with valosin-containing protein (VCP) to activate NF-κB signaling in psoriasis^26^. Beyond keratinocytes, FABP5 is abundantly expressed in skin macrophages, where it mediates DIO-related inflammatory responses in skin lesions^27^. Given FABP5’s emerging role in immunoregulation^28^, its involvement in dietary FA metabolism and signaling within skin macrophages warrants further investigation to elucidate its potential role in obesity-associated psoriasis.

In this study, we investigated the impact of different dietary FAs on obesity and psoriasis using customized HFDs enriched in SFAs (cocoa butter), MUFAs (olive oil), PUFAs (safflower oil), or omega-3 FAs (fish oil). Although all HFDs induced similar levels of obesity, only the cocoa butter HFD accelerated the progression of imiquimod (IMQ)-induced psoriasis. Mechanistically, we found that FABP5 mediates SFA-induced inflammatory IL-1β/IL-17A signaling by coordinating crosstalk between skin-resident macrophages and other immune cells. These findings underscore the critical role of FABP5 in linking dietary fats to psoriasis and uncover how SFAs specifically exacerbate psoriasis through FABP5-mediated mechanisms in the context of obesity.

## Results

### Saturated HFD accelerates IMQ-induced psoriasis progression

To determine the impact of different HFDs on obesity, immunoregulation, and psoriasis development, we developed four types of custom-made HFDs (45% fat), including cocoa butter (SFAs), olive oil (18:1 MUFAs), safflower oil (18:2 PUFAs) or fish oil (n-3 PUFAs), respectively. Low-fat diet (LFD, 5% fat) was used as a control (Figure 1A). All four HFDs shared identical base ingredients except the FA compositions, thus avoiding confounding effects from other dietary components. Of note, cocoa butter primarily contains C16 and C18 SFAs (61%), while olive oil, safflower oil and fish oil are mainly composed of C18:1 MUFAs (71%), C18:2 PUFAs (80%), and n-3 PUFAs (39%), respectively (Figure 1B). Interestingly, compared to LFD-fed mice, mice fed these well-controlled HFD diets exhibited comparable increases in body weight (Figure 1C), fat content (Figure 1D) and body fat percentage (Figure 1E), with unchanged lean mass (Figure S1A, S1B). These suggest that all HFDs induced a similar degree of murine obesity, irrespective of the fat source.

**Figure 1.**
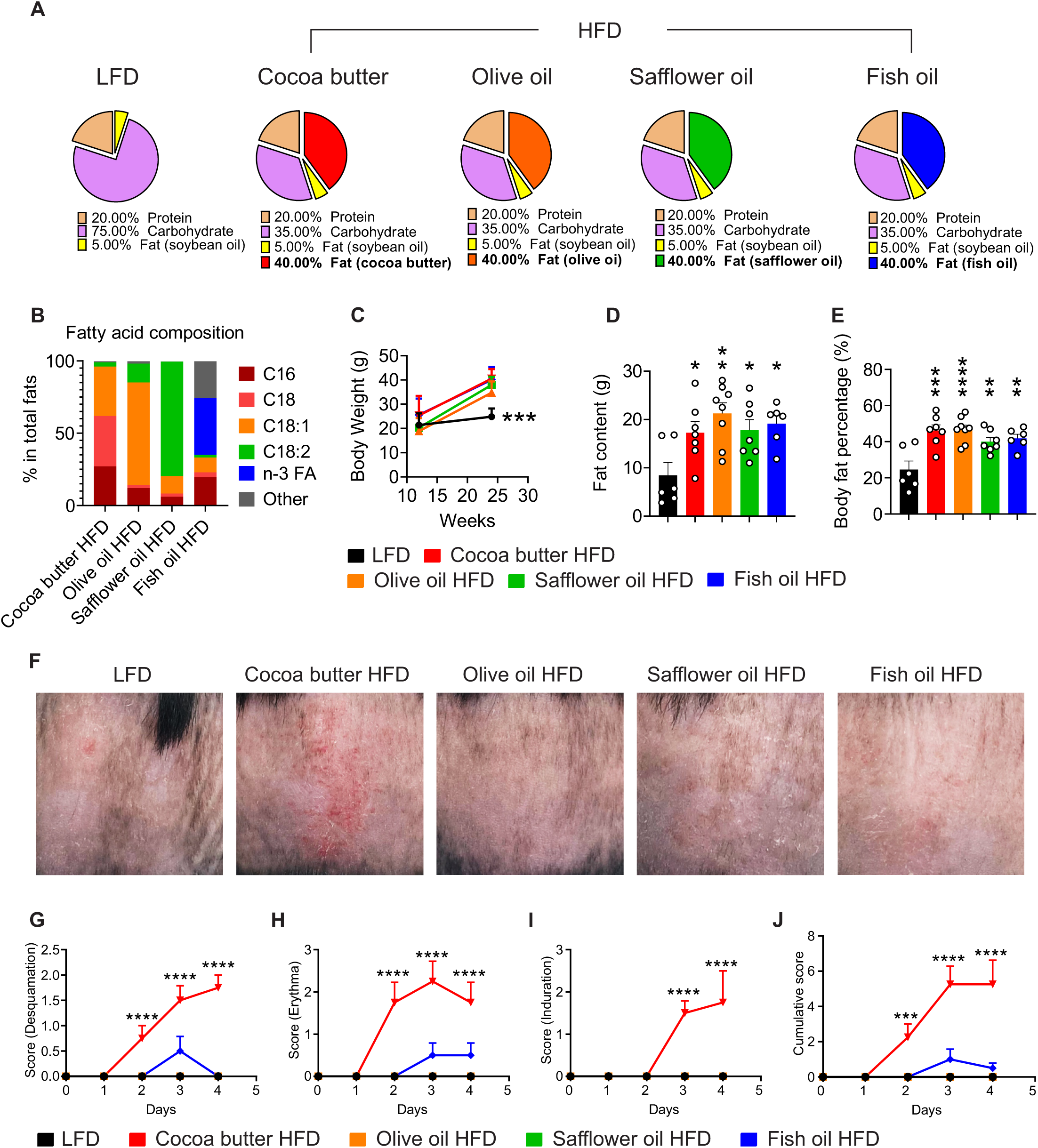
Saturated HFD accelerates IMQ-induced psoriasis progression. A. Compositions of custom-made high-fat diets (HFD) and low-fat diets (LFD). B. Fatty acid (FA) compositions in the fat content of custom-made HFDs. C. Body weight changes of mice fed customized HFDs versus the LFD for 6 months (n=10). D-E. NMR scanning for body fat content (D) and body fat percentage (E) of mice fed customized HFDs versus the LFD. F. Representative images of skin from mice fed customized HFDs or LFD for 6 months followed by 4 days of IMQ treatment (4 mg/mice). G-J. Evaluation of erythema (G), induration (H), and desquamation (I) from day one of IMQ treatment in mice on customized HFDs versus LFD, and the cumulative score (J) summing from G-I. Data are pooled from two independent experiments in panels C-E and G-J, with 4-5 mice per group in each experiment. Data are shown as mean ± SEM, ∗p ≤ 0.05 ∗∗p ≤ 0.01, ∗∗∗p ≤ 0.001, ∗∗∗∗p ≤ 0.0001, one-way ANOVA with Bonferroni’s multiple comparisons test. (See also Figure S1).

To assess the impact of various HFDs on immunoregulation, we analyzed blood immune cell profiles in mice subjected to different dietary conditions. Compared to the LFD, the cocoa butter HFD led to a significant increase in γδ T cells (Figure S1C), while no significant changes were observed in other immune cell populations, including natural killer (NK) cells, CD4^+^ and CD8^+^ T cells, B cells, neutrophils, macrophages and dendritic cells (DCs) (Figure S1D-S1J). In contrast, olive oil and safflower oil HFDs showed minimal effects on peripheral immune populations (Figure S1C-S1J). Mice on the fish oil HFD exhibited a decreased ratio of CD4^+^ and CD8^+^ T cells (Figure S1E, S1F), alongside an increase in neutrophils (Figure S1H). These findings suggest that different HFDs exert distinct influences on peripheral immunoregulation, although the functional implications of these changes remain to be elucidated.

To further explore whether HFD-induced immunoregulation influences susceptibility to immune-mediated psoriasis, we applied imiquimod (IMQ) to the backs of mice fed different HFDs. Interestingly, only mice on the cocoa butter HFD, but not those on the LFD or other HFDs, exhibited a significantly accelerated onset of IMQ-induced psoriasis (Figure 1F). Psoriatic symptoms, evaluated by desquamation (Figure 1G), erythema (Figure 1H), induration (Figure 1I), and cumulative score (Figure 1J), were markedly severer in mice fed the cocoa butter HFD compared to these on the LFD and other HFDs. Collectively, these findings suggest that HFDs rich in SFAs, but not UFAs, exacerbate obesity-associated psoriasis.

### Saturated HFD promotes the infiltration of skin macrophage, TCR γδ^dim^ cell, and neutrophils in IMQ-induced psoriasis

To investigate how saturated HFD exacerbated psoriatic symptoms, we isolated single-cell suspensions from the lesional skin epidermis and dermis of IMQ-treated mice as previously described^29^. Using multi-color flow cytometry, we analyzed the immunophenotype of CD45^+^/Zombie^-^ live immune cells according to a defined gating strategy (Figure S2A). To minimize bias from traditional two-dimensional manual gating, we employed uniform manifold approximation and projection (UMAP) for high-dimensional data visualization across samples including epidermis and dermis from LFD and cocoa butter HFD (Figure 2A-2C). Our analysis demonstrated a marked infiltration of proinflammatory immune cells in the psoriatic skin of mice (Figure 2A, 2B), specifically γδ T cells (populations 0 and 8), F4/80^+^ macrophages (populations 3, 4, 5, 6, 7, 9, and 10), and Ly6G^+^ neutrophils (population 11). In contrast, other types of immune cells, including CD4^+^ T cells, CD8^+^ T cells, DCs (CD11c^+^), and NK cells (NK1.1^+^), were present in lower numbers (Figure 2C).

**Figure 2.**
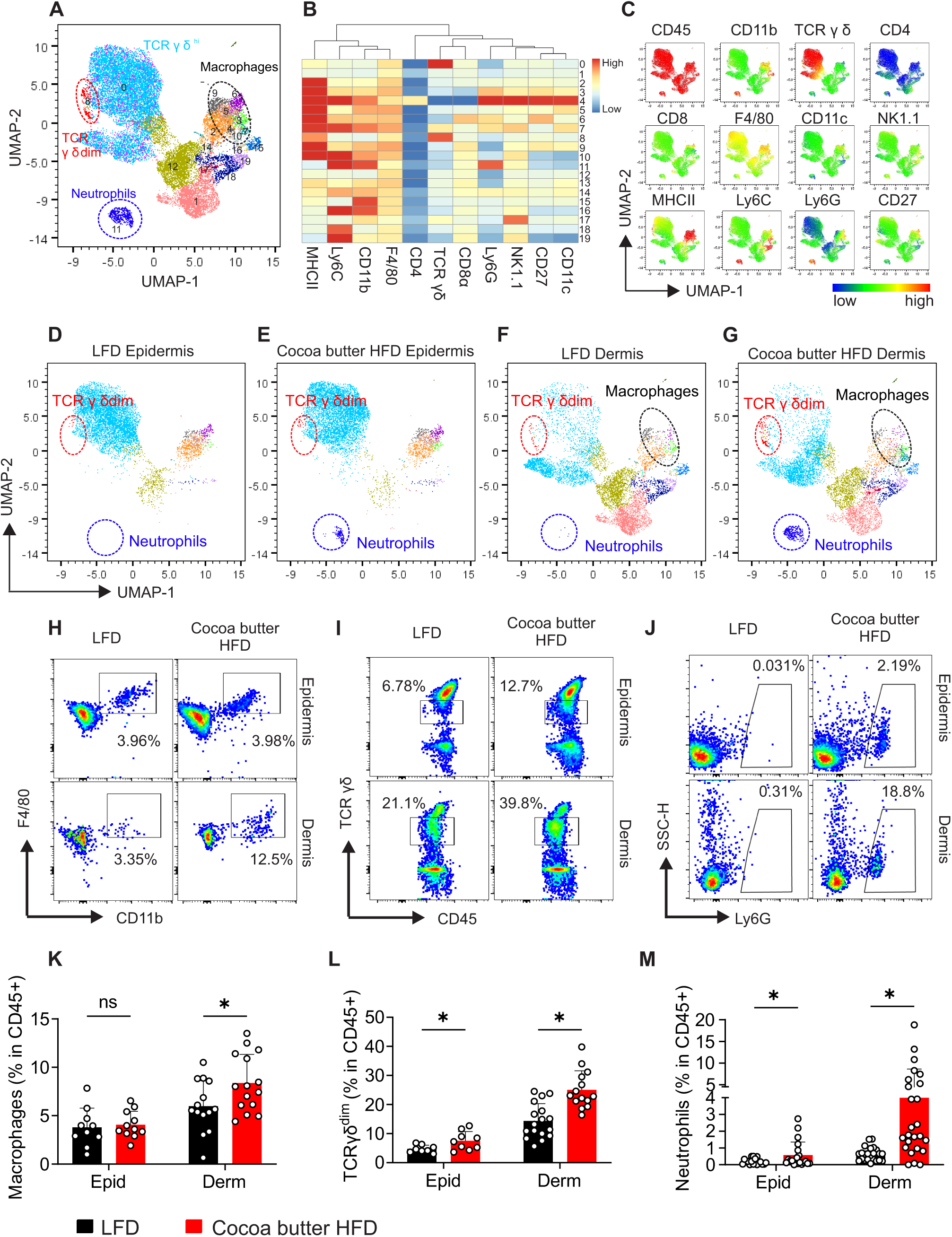
Saturated HFD promotes the infiltration of skin macrophage, TCR γδdim cell, and neutrophils in IMQ-induced psoriasis. A-C. Uniform manifold approximation and projection (UMAP) analysis of representative surface flow staining of skin immune cells following IMQ treatment for 4 days. (A) visualization of skin immune cell clusters, analyzed by FlowSOM. (B) Heatmap displaying indicated markers across 20 cell populations. (C) Individual surface marker signatures. D-G. UMAP visualization of surface staining signatures of dermal and epidermal skin immune cells from mice fed cocoa butter HFD or LFD, treated with IMQ for 4 days. H-J. Representative flow cytometric gating for CD11b+F4/80+ macrophages (H), TCR γδ dim cells (I), and Ly6G+ neutrophils (J) under gate CD45+Zombie-live immune cells in the psoriatic epidermis and dermis from mice on cocoa butter HFD or LFD. K-M. Statistical analysis of ratios of macrophages (K), TCR γδ dim cells (L), and neutrophils (M) in CD45+Zombie-live immune cells in the epidermis (Epid) and dermis (Derm) from mice fed cocoa butter HFD or LFD with IMQ treatment for 4 days. Combination of both dermis and epidermis from 3-4 mice from both LFD and cocoa butter HFD groups was included in panel A-C. Data are pooled from 2-3 independent experiments in panels K-M with 4-5 mice per group. Data are shown as mean ± SEM, ∗p ≤ 0.05, ns, non-significant, unpaired two-tailed multiple t-test. (See also Figure S2).

Given the differences in immune cell compositions between the epidermal and dermal layers, we further analyzed their distribution in psoriatic skin of mice fed either the LFD or the cocoa butter HFD. UMAP analysis showed that the cocoa butter HFD induced infiltration of TCRγδ^dim^ cells and neutrophils in both epidermis and dermis, and macrophages primarily in dermis, compared to the LFD (Figure 2D-2G). Quantitative flow cytometry analysis, following the predefined gating strategy (Figure S2A), confirmed a significant increase in macrophages in the dermis and TCRγδ^dim^ and neutrophils in both skin layers in the cocoa butter HFD group (Figure 2H-2M). In contrast, other immune populations, including CD4^+^ T cells, CD8^+^ T cells, NK cells, B cells, DCs and total γδ T cells, remained unchanged in both skin layers and draining lymph nodes (dLNs) between the cocoa butter HFD and LFD (Figure S2B-S2D), indicating that cocoa butter HFD selectively affects macrophages, TCRγδ^dim^ cells, and neutrophils in psoriatic skin. Notably, other HFDs, including olive oil, safflower oil or fish oil, did not induce significant infiltration of these cell populations compared to the LFD (Figure S2E-S2J). Collectively, our results demonstrated that HFDs enriched in SFAs, but not UFAs, intensified psoriatic inflammation by increasing the infiltration of macrophages, TCRγδ^dim^ cells, and neutrophils in the skin.

### Saturated HFD amplifies IL-1β/IL-17A signaling in IMQ-induced psoriasis

To investigate the function of immune cells induced by the saturated HFD in psoriasis, we first analyzed cytokine expression profiles in the epidermis and dermis across all psoriatic skin samples using UMAP analysis (Figure S3A-S3D). The most abundant expressed cytokines in both layers were IL-17A, IL-1β, TNFα, and IFNγ. Notably, IL-17A was primarily produced by TCR γδ^dim^ cells while IL-1β was mainly produced by macrophages (CD11b^+^F4/80^+^). TNFα and IFNγ appeared to be derived from multiple immune cells (Figure S3E).

We next examined whether the cocoa butter HFD specifically influenced cytokine expression in psoriasis. UMAP analysis showed that cocoa butter HFD increased the population of IL-17A^+^ γδ T cells and IL-1β^+^ macrophages in both the epidermis and dermis of psoriatic skin compared to the LFD, with a corresponding reduction in IL-17A^-^ γδ T cells (Figure 3A-3D). Quantitative flow cytometry analysis confirmed a significant increase in the production of IL-17A^+^ (Figure 3E,3F) and IL-1β^+^ (Figure 3G, 3H) immune cells in the epidermis, dermis, and skin dLNs in the cocoa butter HFD group. In contrast, other immune-derived cytokines, including IFNγ, IL-4, IL-10, IL-23 and TNFα, in the epidermis, dermis and dLN, were unaffected by the cocoa butter HFD (Figure S3F-S3H), highlighting a unique role of the saturated HFD in promoting IL-17A and IL-1β production during psoriasis progression. Consistent with the UMAP analysis, IL-17A and IL-1β induced by the cocoa butter HFD were highly produced by γδ T cells (Figure 3I, 3J) and macrophages (Figure 3K, 3L), respectively. Further analysis demonstrated that IL-17A producing γδ T cells were primarily TCRγδ^dim^ subsets, rather than TCRγδ^high^ cells (Figure S3I-S3L), supporting a critical role of the saturated HFD in activating the function of IL-1β^+^ macrophages and IL-17A^+^ TCR γδ^dim^ T cells in psoriasis development.

**Figure 3.**
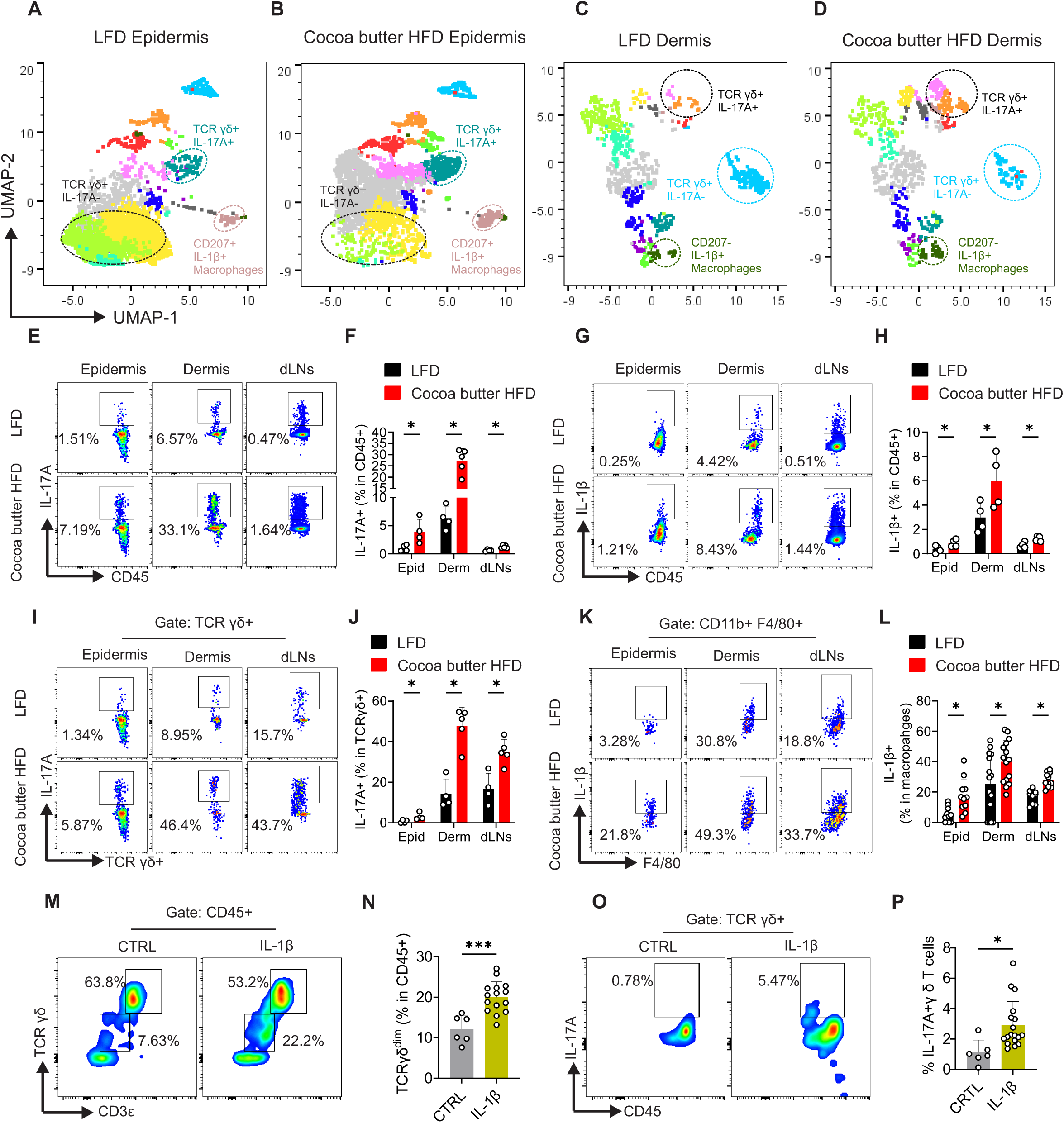
Saturated HFD amplifies IL-1β/IL-17A signaling in IMQ-induced psoriasis. A-D. UMAP visualization of surface and intracellular cytokine staining signature of dermal and epidermal skin immune cells from mice fed cocoa butter HFD or LFD with IMQ treatment for 4 days. E-F. Representative flow cytometric gating for IL-17A+ cells under gate of CD45+Zombie-(live immune cells) in the psoriatic epidermis (Epid), dermis (Derm), and draining lymph nodes (dLNs) from mice on cocoa butter HFD or LFD (E). Statistical analysis was shown in (F). G-H. Statistical analysis of IL-1β+ live immune cell ratio in epidermis, dermis, and dLNs from mice fed cocoa butter HFD or LFD with IMQ treatment for 4 days (G). Statistical analysis was shown in (H). I-J. Statistical analysis of IL-17A+ ratio in γ δ T cells in epidermis, dermis, and dLNs from mice fed cocoa butter HFD or LFD with IMQ treatment for 4 days (I). Statistical analysis was shown in (J). K-L. Statistical analysis of IL-1β+ ratio in CD11b+F4/80+ macrophages in epidermis, dermis, and dLNs from mice fed cocoa butter HFD or LFD with IMQ treatment for 4 days (K). Statistical analysis was shown in (L). M-O. A mixture of epidermal and dermal single-cell suspensions isolated from normal C57BL/6 mice was treated overnight with either 20ng/ml IL-1β or PBS. Surface staining for TCR γδ dim cells and intracellular staining for IL-17A in γ δ T cells is shown in (K). Statistical analysis of TCR γδ dim cell ratio (L) and IL-17A+ ratio in γ δ T cells (M) are displayed. Combination of both dermis and epidermis from 3-4 mice in both LFD and cocoa butter HFD groups was included in panels A-D. Data are from one representative experiment in three independent experiments in panels F, H and J. Data are shown as mean ± SEM, ∗p ≤ 0.05, ∗∗∗p ≤ 0.001, unpaired two-tailed multiple t-test for panels F, H, J, and L, or unpaired two-tailed Student’s t-test for panels N-O. (See also Figure S3).

Prior studies suggested that IL-1β may act as an upstream signal for IL-17A production in γδ T cells in mouse skin^30^. However, whether saturated fats directly induced IL-17A expression in skin γδ T cells remains unclear. To this end, we treated a mixture of single skin cells with IL-1β, bovine serum albumin (BSA)-conjugated palmitic acid (PA), stearic acid (SA), or a BSA control overnight. We found that IL-1β significantly increased TCR γδ^dim^ cell expansion (Figure 3M, 3N) and IL-17A production (Figure 3O, 3P) in skin γδ T cells. However, PA, SA, or BSA did not have the similar effects (Figure S3M-S3O). These results suggest that saturated fats likely promote IL-17A expression in skin γδ T cells indirectly by inducing IL-1β production in skin macrophages.

### Saturated FAs promote IL-1β production in macrophages by inducing extracellular ATP release

To further elucidate how saturated fats enhance IL-1β production in macrophages, we cultured various types of macrophages, including Raw264.7 cell line, mouse peritoneal macrophages (pMac), and GM-CSF-driven bone marrow-derived macrophages (GM-BMMs). These macrophages were treated with SA, OA, docosapentaenoic acid (DPA), or BSA, respectively, and the levels of IL-1β in the culture supernatants were subsequently assessed. Compared to BSA control, SA significantly increased IL-1β secretion across all macrophage types, while OA or DPA did not (Figure 4A-4C). These findings underscore the specific role of SFAs in inducing IL-1β production in macrophages, as opposed to UFAs.

**Figure 4.**
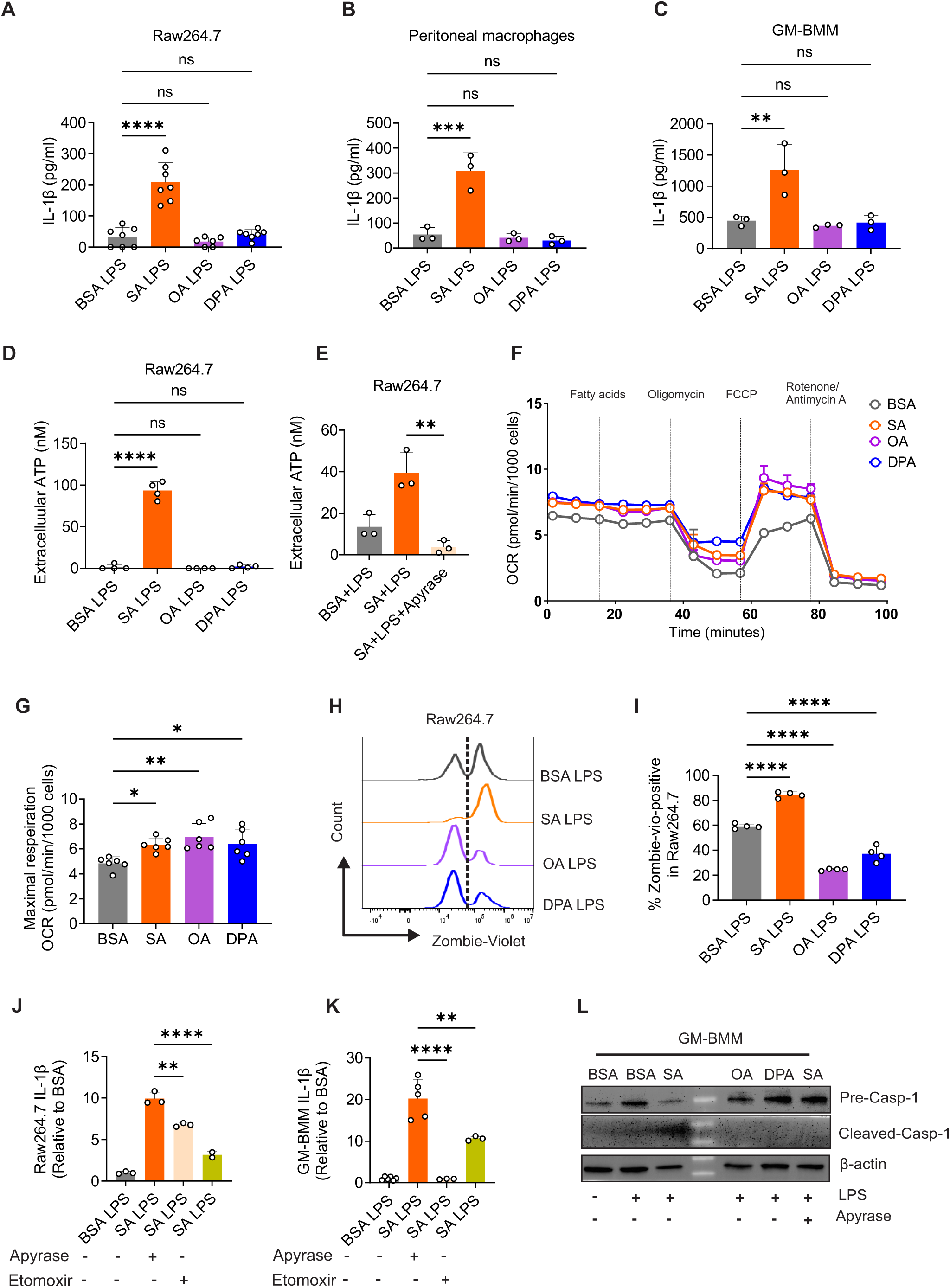
SFAs promote IL-1β production in macrophages by inducing extracellular ATP release. A-C. Supernatant IL-1β concentrations by ELISA of treatments of FAs or BSA on Raw264.7 cells (A) overnight or peritoneal macrophages (B) and GM-BMMs (C) for 6 hours in the presence of LPS. D. Supernatant ATP concentration of Raw264.7 cells treated with FAs or BSA in the presence of LPS for 6 hours. E. Supernatant ATP concentration of Raw264.7 cells treated with SA or BSA in the presence of LPS for 6 hours, with or without 5 units/ml Apyrase. F-G Oxygen consumption rate (OCR) of Raw264.7 cells was measured using a Seahorse analyzer after injection of the indicated drugs at specific time points (F), with statistical analysis of maximal respiration OCR shown in (G). H-I. Raw264.7 cells were treated with FAs or BSA in the presence of LPS for 12 hours. Cell membrane disruption was assessed using Zombie-violet staining and analyzed by flow cytometry (H). Statistical analysis of Zombie-violet+ cells in each group are shown in (I). J-K. treatments of SA or BSA on Raw264.7 cells (J) overnight or GM-BMMs (K) for 6 in the presence of LPS, with (+) or without (-) 5 units/ml Apyrase or 200µM Etomoxir. L. Western blot analysis of pre-caspase-1and cleaved caspase-1(p22) protein levels in cell lysates of GM-BMMs treated with FAs or BSA in the presence or absence of LPS, with or without Apyrase, for 6 hours. Data are shown as mean ± SEM ∗p ≤ 0.05 ∗∗p ≤ 0.01, ∗∗∗p ≤ 0.001, ∗∗∗∗p ≤ 0.0001, ns, non-significant, one-way ANOVA with Bonferroni’s multiple comparisons test. (See also Figure S4).

Extracellular ATP is known to function as a danger signal that triggers inflammasome activation and IL-1β secretion in macrophages^31^. Given that ATP is produced during fatty acid oxidation (FAO) in macrophages^32^, we speculated that SFAs might promote macrophage IL-1β secretion by enhancing ATP release. Indeed, we observed that SA significantly elevated extracellular ATP levels in the supernatant of Raw264.7 cells, regardless of the presence of LPS, while OA, DPA, and BSA did not elicit a similar response (Figure 4D, S4A). Notably, this effect was reversible with the addition of Apyrase, an enzyme that degrades ATP (Figure 4E).

To evaluate whether the increased extracellular ATP by SA correlated with elevated FAO levels in macrophages, we conducted a Seahorse assay on Raw264.7 cells treated with SA, OA, DPA or BSA, respectively. We found that all these FAs increased the oxygen consumption rate (OCR) similarly compared to BSA (Figure 4F, 4G), suggesting that factors beyond FAO might contribute to ATP release from macrophages upon SFA treatment. Interestingly, SA, unlike OA or DPA, increased cell membrane permeability in Raw264.7 cells, as indicated by Zombie-violet staining, both in the presence and absence of LPS (Figure 4H, 4I, Figure S4B). This disruption of cell membrane integrity was also observed in pMac and GM-BMMs (Figure S4C-S4F), reinforcing the notion that SFAs, rather than UFAs, facilitated extracellular ATP release through cell membrane disruption.

To determine if the ATP release induced by SA contributes to IL-1β secretion, we treated the Raw264.7 cells and GM-BMMs with either BSA or SA in the presence of LPS, along with Apyrase or Etomoxir, a FAO inhibitor. Both Apyrase and Etomoxir significantly reduced IL-1β secretion in Raw264.7 cells and GM-BMMs (Figure 4J-4K). Importantly, neither Apyrase nor Etomoxir affected the percentage of Zombie-violet^+^ macrophages treated with SA (Figure S4G, S4H), suggesting that the reduction in IL-1β by these agents was not due to changes in cell membrane permeability.

Additionally, Western blotting analysis demonstrated that SA, not OA or DPA treatment, increased the cleaved form of caspase-1 in cell lysates of GM-BMMs in the presence of LPS, but this effect was negated with the addition of Apyrase (Figure 4L), suggesting SA/ATP-induced caspase-1 activation. Moreover, SA treatment did not affect mRNA levels of IL-1β in either Raw264.7 or GM-BMMs (Figure S4I-S4J), suggesting that SA treatment did not influence IL-1β transcription but promoted IL-1β secretion in macrophages. Altogether, these results demonstrated that SFAs, not UFAs, enhance IL-1β production in macrophages through the induction of extracellular ATP release, thereby contributing to inflammatory processes observed in psoriasis.

### FABP5 mediates saturated FA-induced IL-1β production in macrophages

Long-chain FAs are inherently insoluble in aqueous environments, necessitating the action of FABPs to enhance solubility and facilitate intracellular transport ^23^. Given our previous findings that FABP5, which was highly expressed in skin tissue, plays a critical role in psoriasis pathogenesis^26^, we hypothesized that FABP5 mediated SFA-induced IL-1β production in macrophages. To test this, we first analyzed skin RNA sequencing data from wild-type (WT) or FABP5-knockout (FABP5^-/-^) mice fed the cocoa butter HFD. As shown in the volcano plot and heatmap (Figure 5A, 5B), genes involved in FAO and ATP production, such as *Cyp2e1, Acaa1b, Cyp1a1, Acox3, Aldh1l1, Elovl5, Scd3, Pparg, Aox1, Acss3, Nnmt, Aldh1a7,* were significantly downregulated in FABP5^-/-^ mice compared to WT mice. KEGG pathway analysis showed that these downregulated genes were enriched in pathways related to FAO, including metabolic, PPAR signaling, and FA metabolism pathways (Figure 5C). Additionally, Gene Ontology analysis (GO Biological Process) confirmed that lipid response and FA metabolism pathways were downregulated in the FABP5^-/-^ group (Figure S5A). These findings underscored the critical role of FABP5 in mediating FAO in the skin tissue of mice fed the saturated HFD.

**Figure 5.**
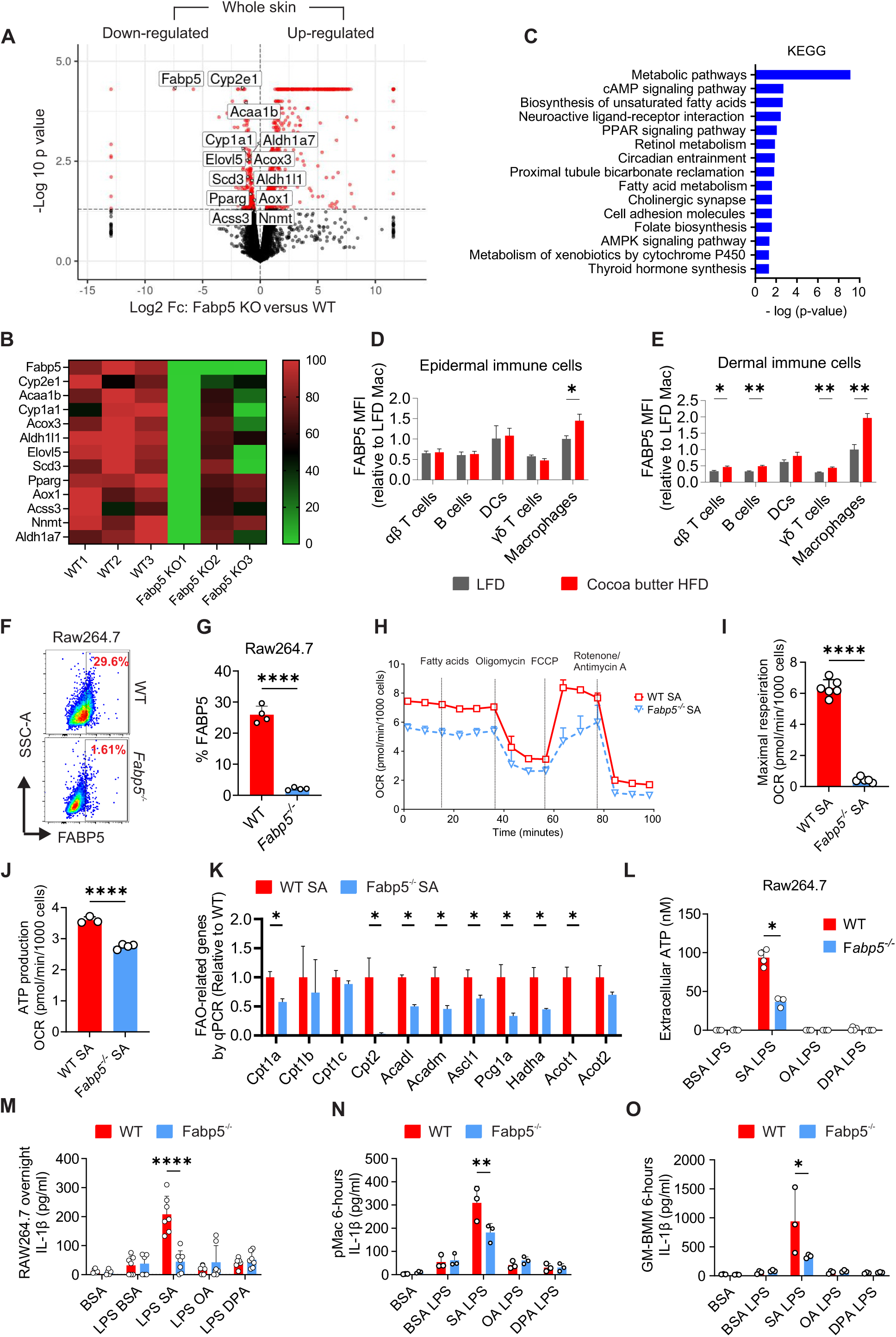
FABP5 is critical in saturated FA-mediated IL-1β production in macrophages. A. The differentially expressed genes (DEGs) of RNA sequencing of the whole skin from wild-type (WT) and Fabp5-/-mice shown in volcano plot with an adjusted p value cutoff of 0.05. Red dots are genes with significantly different expressions, and gray dots are genes with no significant difference. B. Heatmap of downregulated genes in the whole skin of Fabp5-/-mice compared WT mice fed cocoa butter HFD. C. Pathway analysis using KEGG database of DEGs with an adjusted p value cutoff of 0.05 and a Log2 fold change cutoff of -0.5. D-E. Flow cytometric analysis of FABP5 expression level in different cell subsets from epidermis (D) and dermis (E) of mice (n=5) fed cocoa butter HFD or LFD with IMQ treatment for 4 days. F-G. Flow cytometric analysis of FABP5 expression in WT and Fabp5-/-Raw264.7 cells. H-I. OCR of WT and Fabp5-/-Raw264.7 cells was measured a Seahorse analyzer after injection of the indicated drugs at specific time points (H), statistical analysis of maximal respiration OCR shown in (I), and ATP production in (J). K. Expression levels of FAO-related genes in WT and Fabp5-/-Raw264.7 cells treated SA for 4 hours, analyzed by real-time PCR. L. Supernatant ATP concentration of WT and Fabp5-/-Raw264.7 cells treated with FAs or BSA for 6 hours in the presence of LPS. M-O. Supernatant IL-1β concentrations by ELISA of SA or BSA treatment on WT or Fabp5-/-Raw264.7 cells (M) overnight or peritoneal macrophages (N) and GM-BMM (O) for 6 hours in the presence of LPS. Data are shown as mean ± SEM, ∗p ≤ 0.05 ∗∗p ≤ 0.01, ∗∗∗∗p ≤ 0.0001, unpaired two-tailed multiple t-test for panels D, E, K, L, M, N, O, or unpaired two-tailed Student’s t-test for panels G, I and J. (See also Figure S5).

Next, we analyzed FABP5 expression profiles in various immune cell populations in mice fed the LFD or cocoa butter HFD. Interestingly, cocoa butter HFD induced the highest upregulation of FABP5 in macrophages in both epidermis (Figure 5D) and dermis (Figure 5E). To test whether FABP5 was pivotal in mediating FAO in macrophages, we generated a stable FABP5 knockout (Fabp5^-/-^) Raw264.7 cell line using CRISPR-Cas9 technology (Figure S5B) and confirmed complete FABP5 deficiency at both mRNA and protein levels (Figure 5F, 5G, S5C). Through Seahorse assays, we found that FABP5 deficiency resulted in significantly reduced OCR (Figure 5H,5I) and ATP production (Figure 5J) in mitochondria, as well as lower expression of FAO-related genes in Raw264.7 cells (Figure 5K). Moreover, we measured extracellular ATP levels in both WT and FABP5^-/-^ Raw264.7 cells and observed that FABP5 deficiency significantly reduced ATP release upon SA treatment, regardless of LPS priming (Figure 5L, S5D). Notably, FABP5 deficiency did not affect Raw264.7 cell membrane integrity in response to SA treatment (Figure S5E), suggesting that the reduced ATP levels observed in Fabp5^-/-^ Raw264.7 cells are not due to cell membrane disruption. Using primary pMac isolated from WT and Fabp5^-/-^ mice, we observed similar reductions of SA-induced OCR, maximal respiration and ATP production when FABP5 was deficiency (Figure S5F-S5H), further corroborating the critical role of FABP5 in mediating FAO and ATP production in macrophages.

Finally, we measured whether FABP5 deficiency reduced IL-1β secretion using various macrophages, including Raw264.7, pMac, and GM-BMMs. In response to different FA treatment, SA-induced IL-1β secretion was significantly reduced by FABP5 deficiency in Raw264.7 cells (Figure 5M), pMac (Figure 5N), and GM-BMMs (Figure 5O). Of note, FABP5 deficiency did not affect inflammasome-related gene expression in both Raw264.7 cells and GM-BMMs treated with SA (Figure S5I, S5J), suggesting that FABP5 is involved in IL-1β secretion rather than transcriptional regulation. Collectively, these results demonstrate that FABP5 is integral to IL-1β production by macrophages through its role in facilitating SFA oxidation in mitochondria.

### FABP5 deficiency attenuates saturated HFD-related psoriasis with reduction of IL-1β/IL-17A signaling

Given the essential role of FABP5 in promoting FAO and SFA-induced IL-1β production in macrophages, we reasoned that FABP5 was a key factor linking saturated HFD-exacerbated psoriasis. To test this hypothesis, we fed WT and FABP5 whole-body knockout (Fabp5^-/-^) mice with cocoa butter HFD for six months, followed by IMQ-induced psoriasis development. Fabp5^-/-^ mice showed similar body weight as WT mice on the cocoa butter HFD (Figure 6A), suggesting that FABP5 deficiency did not influence saturated HFD-induced murine obesity. Remarkably, FABP5 deficiency significantly alleviated the severity of IMQ-psoriasis symptoms in mice on the cocoa butter HFD (Figure 6B,6C). Flow cytometry analysis demonstrated that FABP5 deficiency significantly reduced neutrophil infiltration in both the epidermis (Figure 6D,6E) and dermis (Figure 6F, 6G) of psoriatic mice. Additionally, TCR γδ^dim^ cells in the epidermis (Figure 6H, 6I) and dermis (Figure 6J, 6K) were also significantly reduced in Fabp5^-/-^ mice compared to WT mice. In contrast, the ratios of other immune cell populations in the epidermis, dermis, and dLNs between WT and Fabp5^-/-^ mice remained the same (Figure S6A-S6C). These *in vivo* studies demonstrated that FABP5 deficiency attenuated saturated HFD-exacerbated psoriasis with reduced infiltration of TCR γδ^dim^ cells and neutrophils.

**Figure 6.**
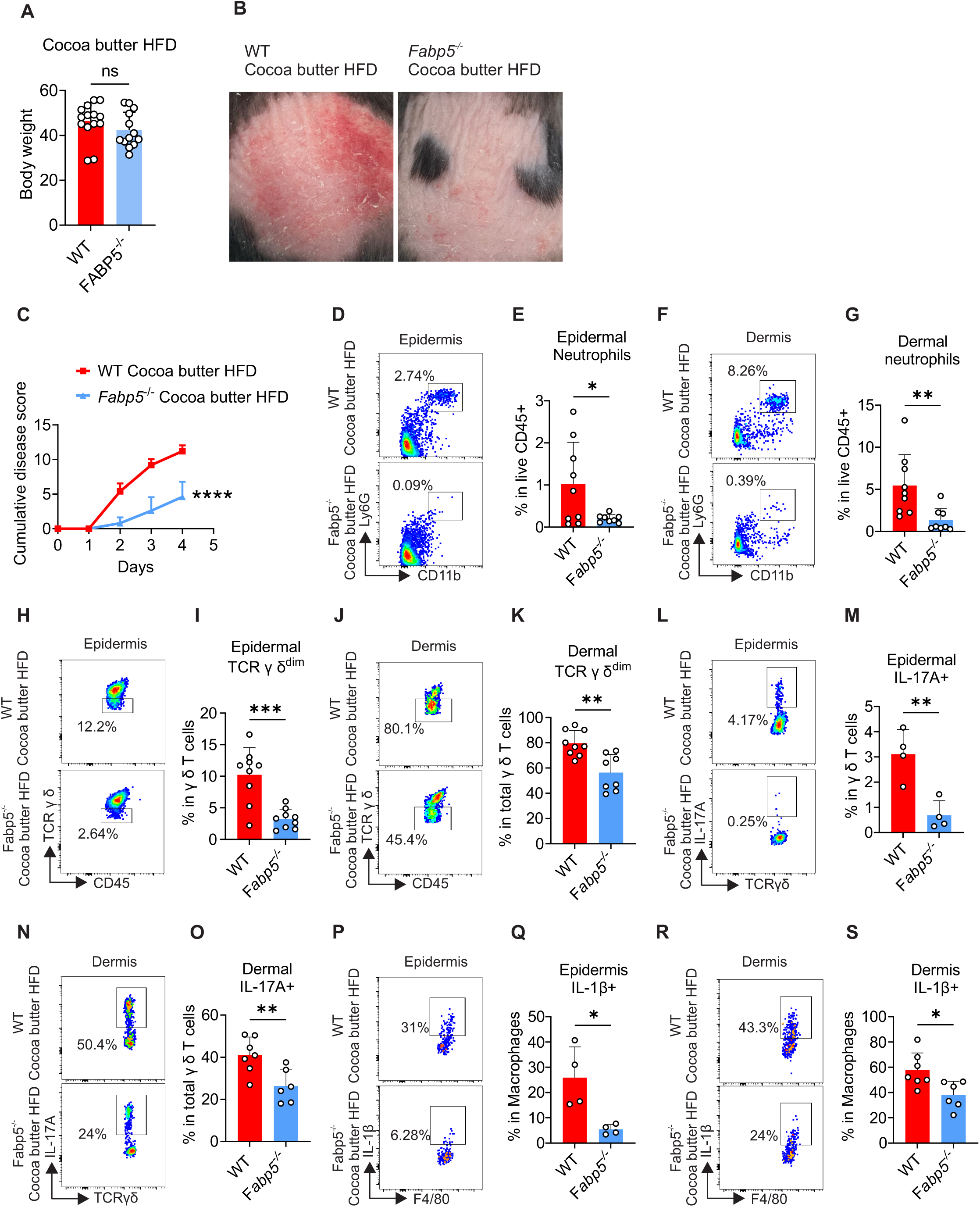
FABP5 deficiency attenuates saturated HFD-related psoriasis with reduction of IL-1β/IL-17A signaling. A. Body weight of WT and Fabp5-/-mice fed cocoa butter HFD for 6 months. B. Representative images of skin from WT and Fabp5-/-mice treated with IMQ for 4 days after being fed cocoa butter HFD or soybean LFD for 6 months. C. The cumulative score was calculated as a sum of the scores of erythema, induration, and desquamation. D-G. Flow cytometric analysis of ratios of CD11b+Ly6G+ neutrophils in the epidermis (D) and dermis (F) of WT and Fabp5-/-mice on cocoa butter HFD, treated with IMQ for 4 days (D). Statistical analysis is shown in (E) and (G). H-K. Flow cytometric analysis of ratios of TCR γ δ dim cells in the epidermis (H) and dermis (J) of WT and Fabp5-/-mice on cocoa butter HFD, treated with IMQ for 4 days. Statistical analysis is shown in (I) and (K). L-O. Flow cytometric analysis of ratios of IL-17A+ in γ δ T cells in the epidermis (L) and dermis (N) of WT and Fabp5-/-mice on cocoa butter HFD, treated with IMQ for 4 days. Statistical analysis is shown in (M) and (O). P-S. Flow cytometric analysis of ratios of IL-1β+ in macrophages in the epidermis (P) and dermis (R) of WT and Fabp5-/-mice on cocoa butter HFD, treated with IMQ for 4 days. Statistical analysis is shown in (Q) and (S). Data are shown as mean ± SEM, ∗p ≤ 0.05 ∗∗p ≤ 0.01, ∗∗∗p ≤ 0.001, ∗∗∗∗p ≤ 0.0001, unpaired two-tailed Student’s t-test. (See also Figure S6).

To further determine how Fabp5^-/-^ mice exhibited reduced TCR γδ^dim^ cells and neutrophils in psoriatic skin tissue, we speculated that FABP5 deficiency reduced saturated HFD-induced IL-1β/IL-17A signaling in skin. Supporting this speculation, we observed a significant decrease of IL-17A^+^ cells in total CD45^+^ immune cells (Figure S6D-S6G) and in γδ T cells in both epidermis (Figure 6L,6M) and dermis (Figure 6N,6O) of FABP5^-/-^ mice compared to WT mice. Moreover, there was a significant reduction in the IL-1β^+^ cells in macrophages in both epidermis (Figure 6P, 6Q) and dermis (Figure 6R-6S). Notably, no significant differences were detected in other proinflammatory cytokines, such as IFNγ, IL-4, in the skin or dLNs between WT and FABP5^-/-^ mice with psoriasis on the cocoa butter HFD (Figure S6H-S6J). Similarly, we found no major differences in the circulating immune cells between these two groups of mice (Figure S6K-S6M). Taken together, these results demonstrated that FABP5 played a crucial role in mediating saturated HFD-related psoriasis by amplifying skin IL-1β/IL-17A signaling in the skin.

### Macrophage-specific FABP5 deficiency attenuates saturated HFD-related psoriasis

To investigate the specific role of FABP5 in skin macrophages in saturated HFD-related psoriasis, we generated macrophage FABP5 conditional knockout mice (Fabp5^f/f^ LysM-Cre) as previously described^26^. To validate the efficiency of FABP5 deficiency in macrophages, we isolated single cells from the skin and peritoneum, followed by flow intracellular staining for FABP5. As anticipated, FABP5 was specifically deficient in CD11b^+^F4/80^+^ macrophages but remained low and unaltered in other immune cell types in the Fabp5^f/f^ LysM-Cre^+^ mice compared to the Fabp5^f/f^ LysM-Cre^-^ mice in the skin (Figure 7A) and peritoneal cells (Figure 7B, S7A).

**Figure 7.**
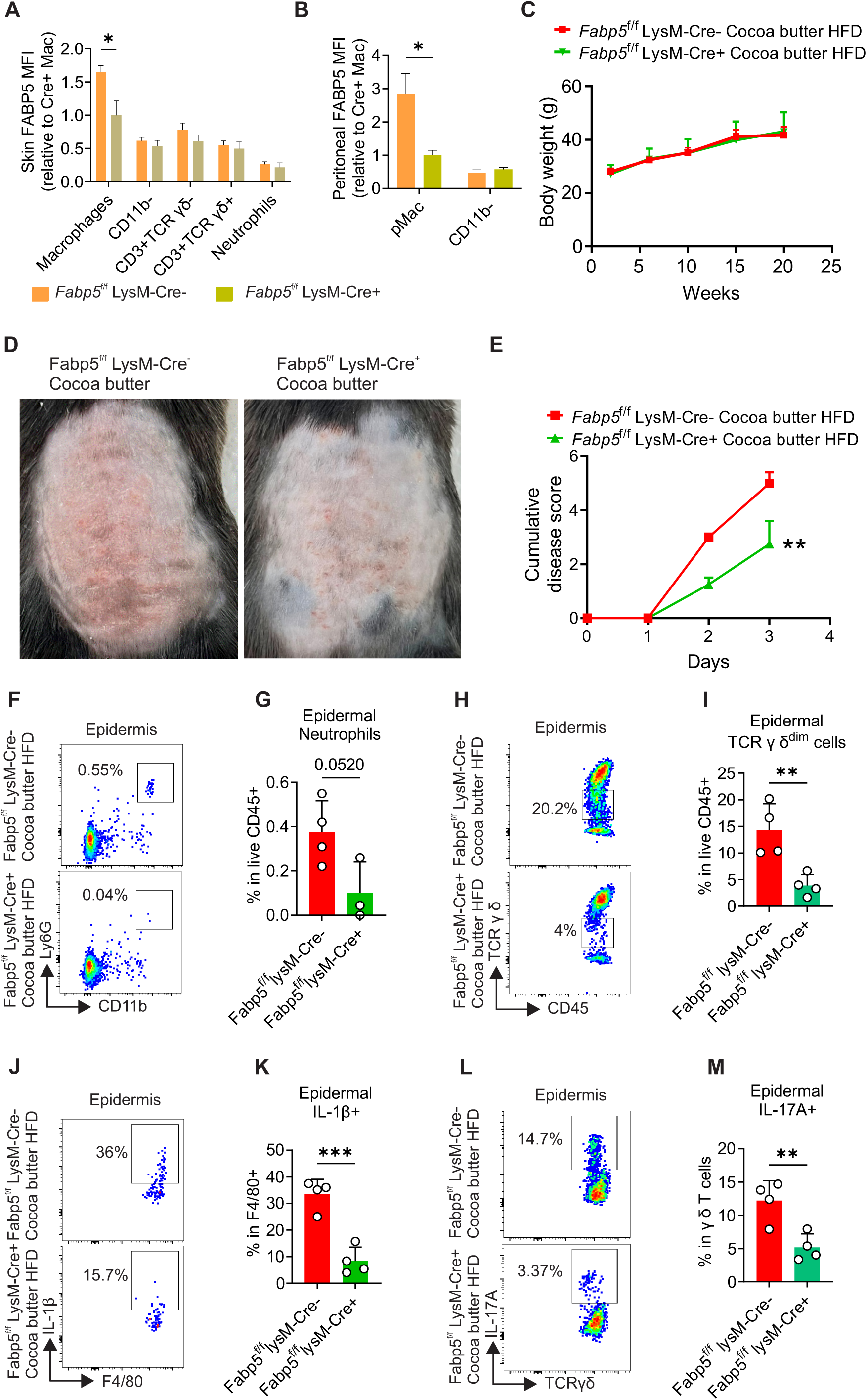
Macrophage-specific FABP5 deficiency attenuates saturated HFD-related psoriasis. A-B. Flow cytometric analysis of FABP5 knockout efficiency by intracellular flow staining for FABP5 expression in different cell subsets from the skin (A) and peritoneum of Fabp5f/f LysM-Cre- and Fabp5f/f LysM-Cre+ mice. C. Body weight curve of Fabp5f/f LysM-Cre- and Fabp5f/f LysM-Cre+ mice fed cocoa butter HFD for 6 months. D-E. Fabp5f/f LysM-Cre- and Fabp5f/f LysM-Cre+ mice were treated with IMQ for 3 days after being fed cocoa butter HFD for 6 months. The representative images of skin (D) and the cumulative disease score (E) was calculated as the sum of erythema, induration, and desquamation scores. F-G. Flow cytometric analysis of ratios of CD11b+Ly6G+ neutrophils in the epidermis of Fabp5f/f LysM-Cre- and Fabp5f/f LysM-Cre+ mice on cocoa butter HFD, treated with IMQ for 3 days (F). Statistical analysis of shown in panel G. H-I. Flow cytometric analysis of ratios of TCR γ δ dim cells in the epidermis of Fabp5f/f LysM-Cre- and Fabp5f/f LysM-Cre+ mice on cocoa butter HFD, treated with IMQ for 4 days (H). Statistical analysis is shown in panel I. J-K. Flow cytometric analysis of ratios of IL-1β+ in F4/80+ macrophages in the epidermis of Fabp5f/f LysM-Cre- and Fabp5f/f LysM-Cre+ mice on cocoa butter HFD, treated with IMQ for 4 days (J). Statistical analysis is shown in panel K. L-M. Flow cytometric analysis of ratios of IL-17A+ γ δ T cells in the epidermis of Fabp5f/f LysM-Cre- and Fabp5f/f LysM-Cre+ mice on cocoa butter HFD, treated with IMQ for 4 days (L). Statistical analysis is shown in panel M. Data are shown as mean ± SEM, ∗p ≤ 0.05, ∗∗p ≤ 0.01, unpaired two-tailed multiple t-test in panels A-B, or unpaired two-tailed Student’s t-test in panels C, E, F, G, H, I, J, K L, and M. (See also Figure S7).

We then fed Fabp5^f/f^ LysM-Cre mice the cocoa butter HFD followed by psoriasis induction with IMQ treatment. Both Fabp5^f/f^ LysM-Cre^-^ and Fabp5^f/f^ LysM-Cre^+^ mice developed similar levels of obesity on the cocoa butter HFD, as indicated by comparable body weights and fat deposition (Figure 7C, S7B, S7C), suggesting that macrophage-specific FABP5 deficiency did not affect the obese status of mice.

Interestingly, consistent with findings in whole-body Fabp5^-/-^ mice, Fabp5^f/f^ LysM-Cre^+^ mice on the cocoa butter HFD exhibited significantly reduced psoriasis symptoms compared to Fabp5^f/f^ LysM-Cre^-^ mice following IMQ treatment (Figure 7D,7E). Similarly, Fabp5^f/f^ LysM-Cre^+^ mice showed a significant reduction in the infiltration of neutrophils (Figure 7F,7G) and TCR γδ^dim^ cells (Figure 7HI,7I) in the epidermis compared to Fabp5^f/f^ LysM-Cre^-^ mice with psoriasis on the cocoa butter HFD. These data suggested that macrophage-specific FABP5 deficiency attenuated psoriasis progression by reducing the infiltration of proinflammatory immune cells.

Moreover, we observed a significant decrease of IL-1β^+^ macrophages (Figure 7J,7K) and IL-17A^+^ γδ T cells (Figure 7L,7M) in the psoriatic epidermis, along with reduced IL-17A^+^ γδ T cells in dLNs (Figure S7D) in Fabp5^f/f^ LysM-Cre^+^ mice compared to Fabp5^f/f^ LysM-Cre^-^ mice on the cocoa butter HFD. Of note, no significant difference in IL-1β production was observed in dermal macrophages in these mice (Figure S7E). This may be due to dermal macrophages in psoriasis being primarily derived from peripheral monocytes^33^, which exhibited lower FABP5 expression compared to skin-resident macrophages^34^, as shown in both humans (Figure S7F, S7G) and mice (Figure S7H). These findings suggested that FABP5 deficiency in skin macrophages dampened IL-1β/IL-17A signaling in saturated HFD-related psoriasis. By contrast, we observed no significant differences in other skin immune cell populations (Figure S7I-S7K) or in the expression of other pro-inflammatory cytokines (Figure S7L-S7N) in the skin and dLNs between the two groups. In addition, there were no major differences in the immunophenotypes in peripheral blood and spleen (Figure S7O-S7P). Collectively, these data demonstrated that FABP5 expression in macrophages played a critical role in linking saturated HFD and psoriasis development.

## Discussion

Psoriasis, the most common immune-driven inflammatory skin disease, affects over 100 million people globally^35^. While genetic predisposition is a significant risk factor, metabolic and environmental influences, such as HFDs, obesity, and lifestyle factors, also play critical roles in exacerbating the incidence and severity of psoriasis ^36^. The increasing prevalence of HFD-induced obesity has contributed to a global rise in psoriasis cases ^37,38^, yet the molecular mechanisms linking dietary lipids to psoriasis risk remain poorly understood.

In this study, we employed custom-designed HFDs with distinct fat compositions to investigate their effects on psoriasis. Our results demonstrated that HFDs high in saturated fats, but not unsaturated fats, exacerbated psoriasis onset and severity in murine models. Mechanistically, SFAs drove FABP5-mediated mitochondrial FAO, extracellular ATP release and IL-1β production in skin macrophages. Deleting FABP5, either globally or specifically in macrophages, significantly reduced IL-1β/IL-17A signaling neutrophil infiltration, and psoriatic inflammation linked to saturated HFDs.

Interestingly, the severity of psoriasis was not linked to obesity itself, as both saturated and unsaturated HFDs induced similar levels of obesity. Rather, the pro-psoriatic effects were specific to SFAs, which amplify the IL-1β/IL-17A signaling axis. Flow cytometry and UMAP analyses showed increased proinflammatory immune cell populations, including macrophages, γδ T cells, and neutrophils in psoriatic skin of mice fed the saturated HFD. Additionally, cytokine analysis demonstrated that SFAs uniquely elevated IL-1β production by macrophages, which in turn enhanced IL-17A expression in γδ T cells and promoted neutrophil recruitment^30,39^ ^40^. *In vitro* studies further confirmed that IL-1β, rather than SFAs themselves, directly induced IL-17A expression in γδ T cells, emphasizing the role of IL-1β as a key intermediary in saturated HFD-induced psoriatic inflammation.

Macrophages, as key responders to lipid stimuli, produce diverse cytokine and chemokine profiles in response to different types of FAs, contributing to the pathogenesis of inflammatory diseases, including psoriasis ^41–43^. In this study, we found that SFAs, not UFAs, induced IL-1β secretion in activated macrophages. While palmitic acid (PA, 16:0) has been implicated in inflammasome activation via protein palmitoylation ^44,45^, our study showed that stearic acid (SA, 18:0) elicited even stronger inflammasome activation and IL-1β secretion. These observations suggest that mechanisms beyond palmitoylation contribute to SFA-induced inflammasome activation.

Further investigation revealed that SFA treatment caused cell membrane disruption and release of extracellular ATP, a known danger signal for inflammasome activation and IL-1β secretion^31,46^. Pharmacological inhibition of ATP generation using etomoxir or degradation of extracellular ATP with Apyrase significantly reduced SFA-induced IL-1β secretion in macrophages, highlighting the critical role of extracellular ATP in linking SFAs to inflammatory responses.

FABP5, an intracellular lipid chaperone, plays a pivotal role in regulating lipid transport and metabolism in skin macrophages and keratinocytes^28,47^. RNA sequencing of skin tissue from WT and Fabp5^-/-^ mice fed the saturated HFD showed that FABP5 deficiency dampened FAO pathways. Intracellular flow staining demonstrated that saturated HFD induced the highest FABP5 expression in skin macrophages, and FABP5 deletion reduced SFA-induced FAO, extracellular ATP levels, and IL-1β secretion. Notably, FABP5 deficiency did not cause cell membrane disruption, suggesting that FABP5 mediates ATP generation rather than facilitating ATP release through membrane disruption. Interestingly, FABP4 has been shown to promote SFA-induced ceramide production and cell death^48^. It is likely that FABP4 and FABP5 may coordinate SFA-induced effects in macrophages.

To further evaluate the role of FABP5 in saturated HFD-related psoriasis, we used both whole-body Fabp5 knockout and macrophage-specific FABP5 conditional knockout mice (LysM-Cre^+^FABP5^fl/fl^). FABP5 deficiency attenuated psoriasis symptoms without affecting obesity levels in saturated HFD-fed mice, suggesting that FABP5 deficiency uncouples obesity from psoriasis-associated inflammation. Flow cytometric analysis demonstrated FABP5 deficiency reduced IL-1β production in langerin (CD207)^+^ epidermal macrophages, highlighting their critical role in SFA-induced FABP5/IL-1β signaling.

Interestingly, CD207^-^ dermal macrophages also exhibited reduced IL-1β production in whole-body FABP5 knockout mice but not in LysM-Cre^+^FABP5^fl/fl^ mice. This discrepancy could stem from differences in FABP5 expression between tissue-resident macrophages and infiltrating monocyte-derived macrophages during psoriatic inflammation^49,50^. In addition, monocytes exhibited lower basal expression of lysozyme than macrophages^51^. Therefore, FABP5 deficiency might be less effective in these immature monocytes when using a lysozyme gene-driven conditional knockout approach. These findings suggest that FABP5 plays a more pronounced role in resident macrophages than in infiltrating monocytes, further underscoring its importance in saturated HFD-induced inflammation.

Although FABP5 is expressed in other immune cells, such as T cells^28^, its levels are significantly lower than in skin macrophages, which may explain the lack of systemic effects upon FABP5 deficiency. Moreover, as a FA chaperone, FABP5 exhibits more pronounced immunoregulatory effects in HFD-induced obese mice than in non-obese lean mice. Our data demonstrated that FABP5 in skin macrophages mediates SFA-induced IL-1β signaling, driving the pathogenesis of saturated HFD-associated psoriasis. These findings align with previous studies identifying FABP5 as a lipid sensor implicated in macrophage-associated inflammatory diseases^4,27^.

In conclusion, our study highlights the critical role of FABP5 in linking saturated HFD intake to psoriasis development by promoting IL-1β-mediated immune responses. Targeting FABP5 activity may therefore represent a promising therapeutic strategy for managing psoriasis, particularly in the context of saturated HFD-induced obesity.

### Limitation of the study

While obesity is a recognized risk factor for psoriasis, the influence of different fat sources in HFD-induced obesity on psoriasis development varies. Our study demonstrated that saturated, but not unsaturated, HFDs exacerbate psoriasis onset. However, it remains unclear whether unsaturated HFDs might play a protective role in psoriasis. We also identified FABP5 as a critical mediator linking between saturated HFDs to psoriasis. Notably, the dual expression of FABP5 in both keratinocytes and immune cells raises questions about its potential roles in keratinocytes and other skin-infiltrating immune cells. Further investigation is needed to determine whether FABP5 expression in these cells is involved in saturated fat-exacerbated psoriasis.

## Acknowledgments

We thank the Department of Pathology at the University of Iowa for the access to use the Cytek Aurora flow cytometry. B.L. thanks the funding support from NIH grants R01AI137324, R01CA180986, and U01CA272424.

## Author contributions

J.Y., J.H. performed most of the experiments and analyzed the data. M.S.Y., X.J., A.A., S.L., Z.W., X.H., and J.S. provided technical assistance. B.L. designed the experiments.

A.J. helped with intellectual inputs and paper writing. B.L. and J.Y. wrote the paper.

## STAR Methods

### RESOURCE AVAILABILITY

#### Lead contact

Further information and requests for reagents may be directed to and will be fulfilled by the Lead Contact, Bing Li.

#### Material availability

All unique/stable reagents generated in this study are available from the lead contact with a completed materials transfer agreement.

### KEY RESOURCE TABLE

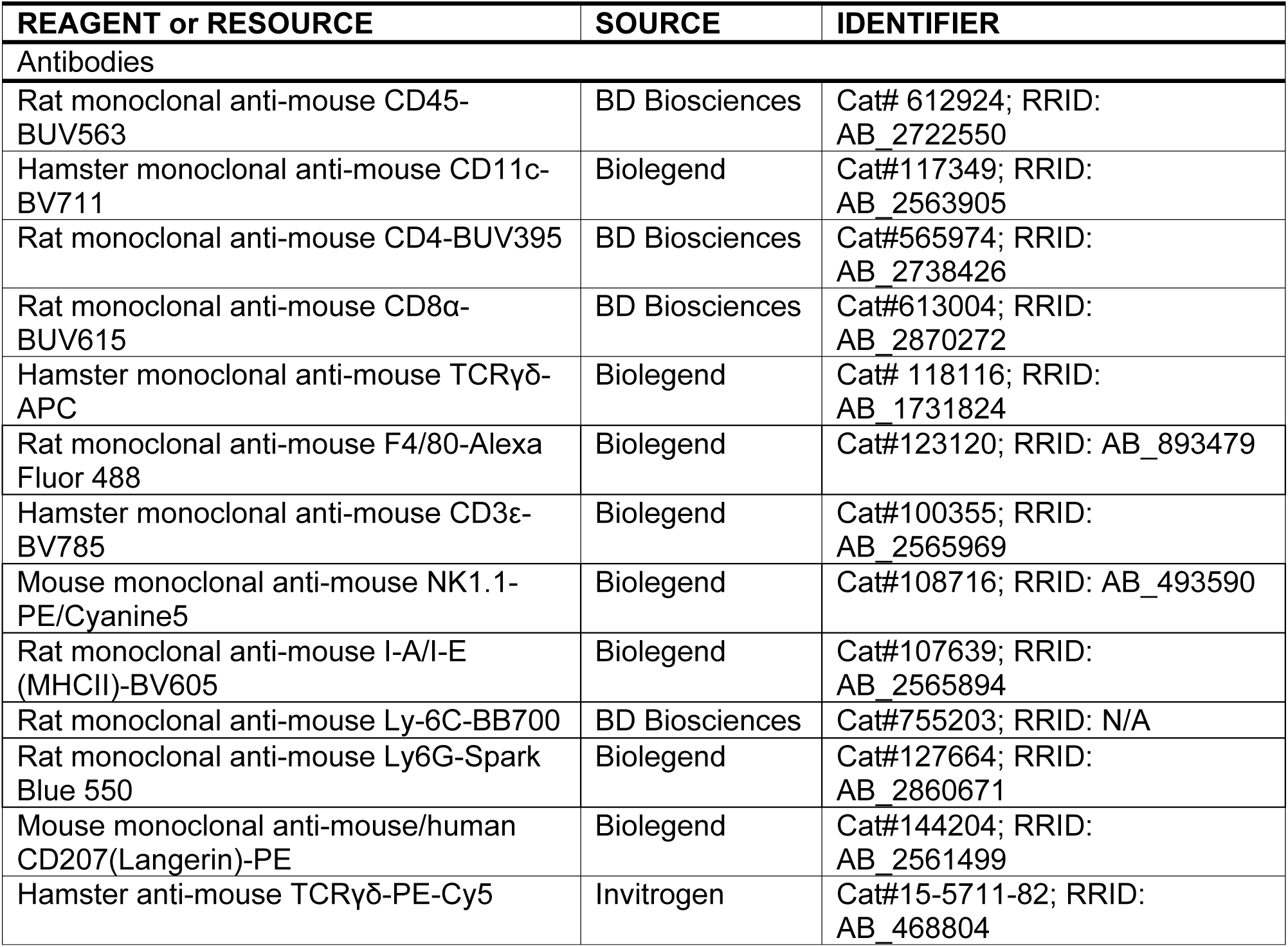
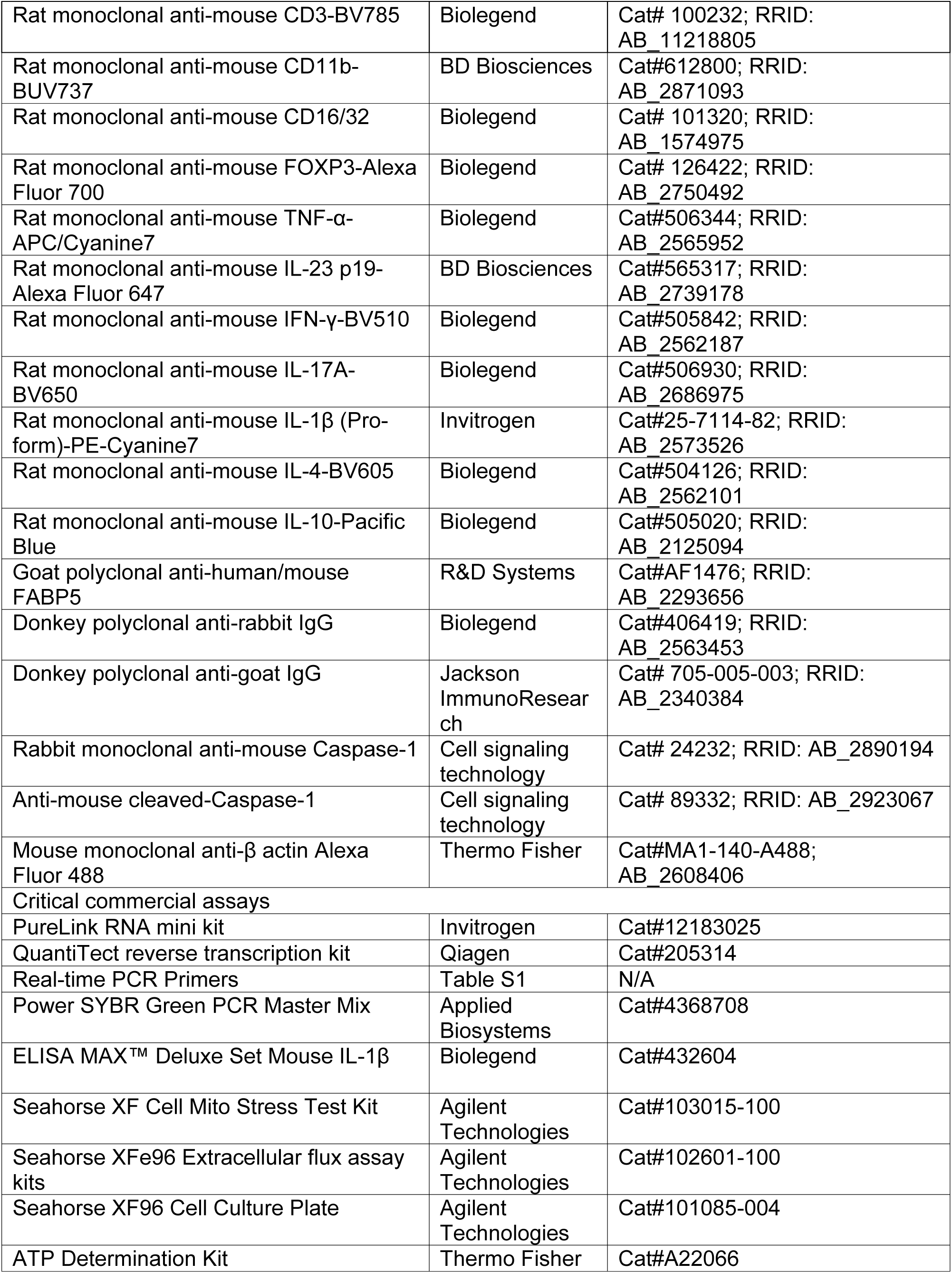
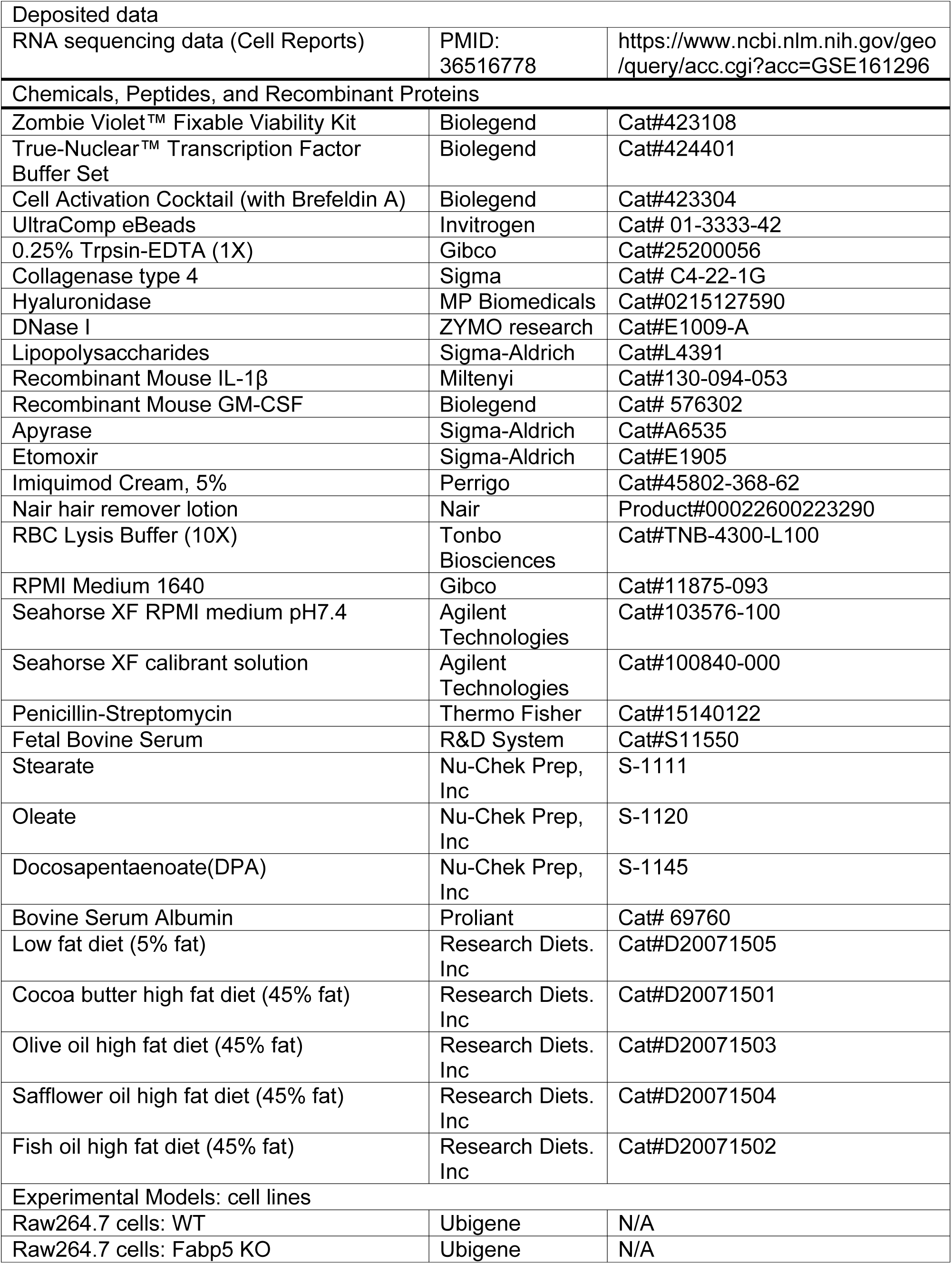
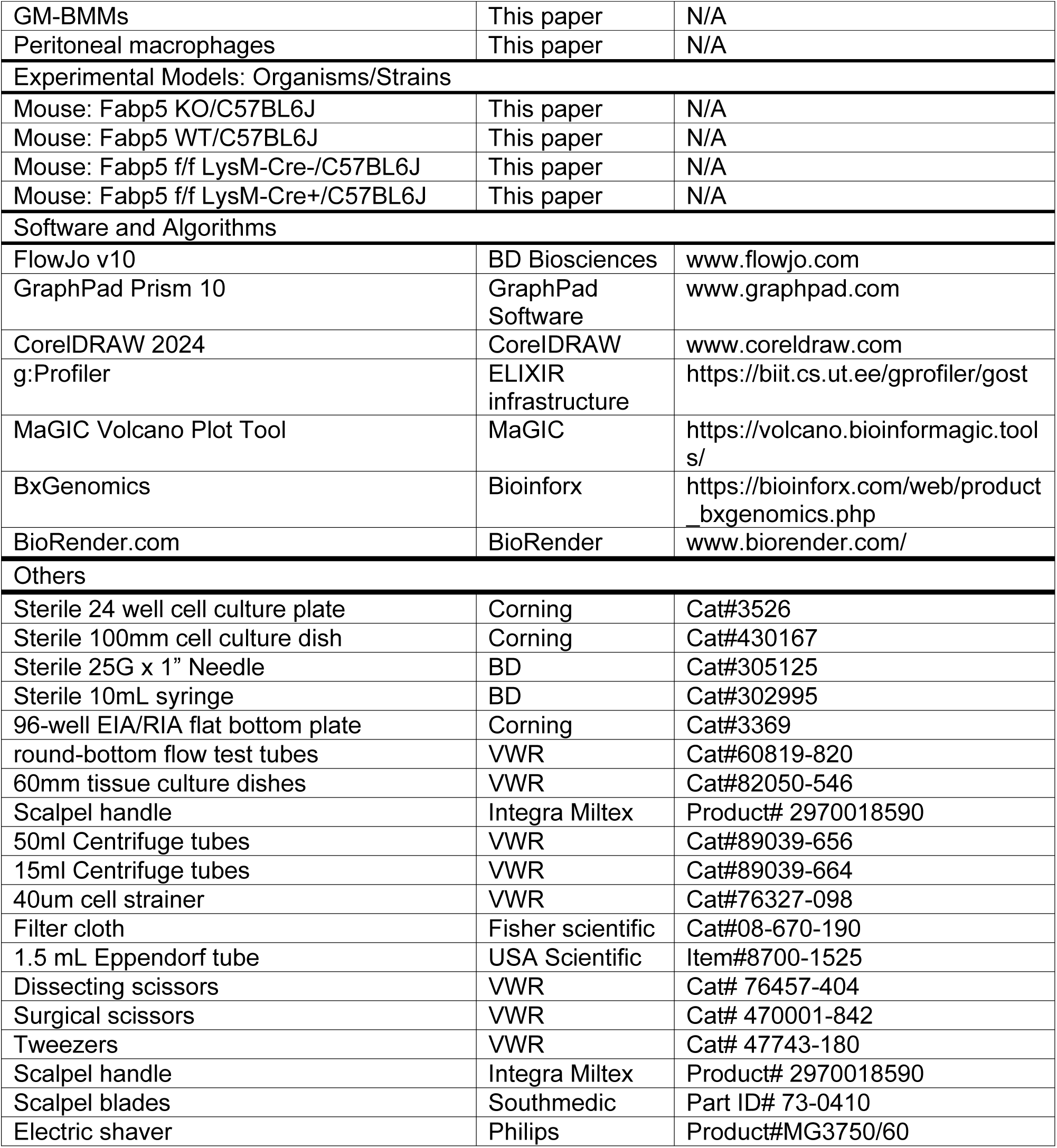

| REAGENT or RESOURCE | SOURCE | IDENTIFIER |
| --- | --- | --- |
| Antibodies |  |  |
| Rat monoclonal anti-mouse CD45-BUV563 | BD Biosciences | Cat# 612924; RRID: AB_2722550 |
| Hamster monoclonal anti-mouse CD11c-BV711 | Biolegend | Cat#117349; RRID: AB_2563905 |
| Rat monoclonal anti-mouse CD4-BUV395 | BD Biosciences | Cat#565974; RRID: AB_2738426 |
| Rat monoclonal anti-mouse CD8α-BUV615 | BD Biosciences | Cat#613004; RRID: AB_2870272 |
| Hamster monoclonal anti-mouse TCR$\gamma\delta$-APC | Biolegend | Cat# 118116; RRID: AB_1731824 |
| Rat monoclonal anti-mouse F4/80-Alexa Fluor 488 | Biolegend | Cat#123120; RRID: AB_893479 |
| Hamster monoclonal anti-mouse CD3ε-BV785 | Biolegend | Cat#100355; RRID: AB_2565969 |
| Mouse monoclonal anti-mouse NK1.1-PE/Cyanine5 | Biolegend | Cat#108716; RRID: AB_493590 |
| Rat monoclonal anti-mouse I-A/I-E (MHCII)-BV605 | Biolegend | Cat#107639; RRID: AB_2565894 |
| Rat monoclonal anti-mouse Ly-6C-BB700 | BD Biosciences | Cat#755203; RRID: N/A |
| Rat monoclonal anti-mouse Ly6G-Spark Blue 550 | Biolegend | Cat#127664; RRID: AB_2860671 |
| Mouse monoclonal anti-mouse/human CD207(Langerin)-PE | Biolegend | Cat#144204; RRID: AB_2561499 |
| Hamster anti-mouse TCR$\gamma\delta$-PE-Cy5 | Invitrogen | Cat#15-5711-82; RRID: AB_468804 |
| Rat monoclonal anti-mouse CD3-BV785 | Biolegend | Cat# 100232; RRID: AB_11218805 |
| Rat monoclonal anti-mouse CD11b-BUV737 | BD Biosciences | Cat#612800; RRID: AB_2871093 |
| Rat monoclonal anti-mouse CD16/32 | Biolegend | Cat# 101320; RRID: AB_1574975 |
| Rat monoclonal anti-mouse FOXP3-Alexa Fluor 700 | Biolegend | Cat# 126422; RRID: AB_2750492 |
| Rat monoclonal anti-mouse TNF-α-APC/Cyanine7 | Biolegend | Cat#506344; RRID: AB_2565952 |
| Rat monoclonal anti-mouse IL-23 p19-Alexa Fluor 647 | BD Biosciences | Cat#565317; RRID: AB_2739178 |
| Rat monoclonal anti-mouse IFN-γ-BV510 | Biolegend | Cat#505842; RRID: AB_2562187 |
| Rat monoclonal anti-mouse IL-17A-BV650 | Biolegend | Cat#506930; RRID: AB_2686975 |
| Rat monoclonal anti-mouse IL-1β (Pro-form)-PE-Cyanine7 | Invitrogen | Cat#25-7114-82; RRID: AB_2573526 |
| Rat monoclonal anti-mouse IL-4-BV605 | Biolegend | Cat#504126; RRID: AB_2562101 |
| Rat monoclonal anti-mouse IL-10-Pacific Blue | Biolegend | Cat#505020; RRID: AB_2125094 |
| Goat polyclonal anti-human/mouse FABP5 | R&D Systems | Cat#AF1476; RRID: AB_2293656 |
| Donkey polyclonal anti-rabbit IgG | Biolegend | Cat#406419; RRID: AB_2563453 |
| Donkey polyclonal anti-goat IgG | Jackson ImmunoResearch | Cat# 705-005-003; RRID: AB_2340384 |
| Rabbit monoclonal anti-mouse Caspase-1 | Cell signaling technology | Cat# 24232; RRID: AB_2890194 |
| Anti-mouse cleaved-Caspase-1 | Cell signaling technology | Cat# 89332; RRID: AB_2923067 |
| Mouse monoclonal anti-β actin Alexa Fluor 488 | Thermo Fisher | Cat#MA1-140-A488; AB_2608406 |
| Critical commercial assays |  |  |
| PureLink RNA mini kit | Invitrogen | Cat#12183025 |
| QuantiTect reverse transcription kit | Qiagen | Cat#205314 |
| Real-time PCR Primers | Table S1 | N/A |
| Power SYBR Green PCR Master Mix | Applied Biosystems | Cat#4368708 |
| ELISA MAX™ Deluxe Set Mouse IL-1β | Biolegend | Cat#432604 |
| Seahorse XF Cell Mito Stress Test Kit | Agilent Technologies | Cat#103015-100 |
| Seahorse XFe96 Extracellular flux assay kits | Agilent Technologies | Cat#102601-100 |
| Seahorse XF96 Cell Culture Plate | Agilent Technologies | Cat#101085-004 |
| ATP Determination Kit | Thermo Fisher | Cat#A22066 |
| Deposited data |  |  |
| RNA sequencing data (Cell Reports) | PMID:<br>36516778 | https://www.ncbi.nlm.nih.gov/geo/query/acc.cgi?acc=GSE161296 |
| Chemicals, Peptides, and Recombinant Proteins |  |  |
| Zombie Violet™ Fixable Viability Kit | Biolegend | Cat#423108 |
| True-Nuclear™ Transcription Factor Buffer Set | Biolegend | Cat#424401 |
| Cell Activation Cocktail (with Brefeldin A) | Biolegend | Cat#423304 |
| UltraComp eBeads | Invitrogen | Cat# 01-3333-42 |
| 0.25% Trpsin-EDTA (1X) | Gibco | Cat#25200056 |
| Collagenase type 4 | Sigma | Cat# C4-22-1G |
| Hyaluronidase | MP Biomedicals | Cat#0215127590 |
| DNase I | ZYMO research | Cat#E1009-A |
| Lipopolysaccharides | Sigma-Aldrich | Cat#L4391 |
| Recombinant Mouse IL-1β | Miltenyi | Cat#130-094-053 |
| Recombinant Mouse GM-CSF | Biolegend | Cat# 576302 |
| Apyrase | Sigma-Aldrich | Cat#A6535 |
| Etomoxir | Sigma-Aldrich | Cat#E1905 |
| Imiquimod Cream, 5% | Perrigo | Cat#45802-368-62 |
| Nair hair remover lotion | Nair | Product#00022600223290 |
| RBC Lysis Buffer (10X) | Tonbo Biosciences | Cat#TNB-4300-L100 |
| RPMI Medium 1640 | Gibco | Cat#11875-093 |
| Seahorse XF RPMI medium pH7.4 | Agilent Technologies | Cat#103576-100 |
| Seahorse XF calibrant solution | Agilent Technologies | Cat#100840-000 |
| Penicillin-Streptomycin | Thermo Fisher | Cat#15140122 |
| Fetal Bovine Serum | R&D System | Cat#S11550 |
| Stearate | Nu-Chek Prep, Inc | S-1111 |
| Oleate | Nu-Chek Prep, Inc | S-1120 |
| Docosapentaenoate(DPA) | Nu-Chek Prep, Inc | S-1145 |
| Bovine Serum Albumin | Proliant | Cat# 69760 |
| Low fat diet (5% fat) | Research Diets. Inc | Cat#D20071505 |
| Cocoa butter high fat diet (45% fat) | Research Diets. Inc | Cat#D20071501 |
| Olive oil high fat diet (45% fat) | Research Diets. Inc | Cat#D20071503 |
| Safflower oil high fat diet (45% fat) | Research Diets. Inc | Cat#D20071504 |
| Fish oil high fat diet (45% fat) | Research Diets. Inc | Cat#D20071502 |
| Experimental Models: cell lines |  |  |
| Raw264.7 cells: WT | Ubigen | N/A |
| Raw264.7 cells: Fabp5 KO | Ubigen | N/A |
| GM-BMMs | This paper | N/A |
| Peritoneal macrophages | This paper | N/A |
| Experimental Models: Organisms/Strains |  |  |
| Mouse: Fabp5 KO/C57BL6J | This paper | N/A |
| Mouse: Fabp5 WT/C57BL6J | This paper | N/A |
| Mouse: Fabp5 f/f LysM-Cre-/C57BL6J | This paper | N/A |
| Mouse: Fabp5 f/f LysM-Cre+/C57BL6J | This paper | N/A |
| Software and Algorithms |  |  |
| FlowJo v10 | BD Biosciences | www.flowjo.com |
| GraphPad Prism 10 | GraphPad Software | www.graphpad.com |
| CorelDRAW 2024 | CorelDRAW | www.coreldraw.com |
| g:Profiler | ELIXIR infrastructure | https://biit.cs.ut.ee/gprofiler/gost |
| MaGIC Volcano Plot Tool | MaGIC | https://volcano.bioinformagic.tools/ |
| BxGenomics | Bioinforx | https://bioinforx.com/web/product_bxgenomics.php |
| BioRender.com | BioRender | www.biorender.com/ |
| Others |  |  |
| Sterile 24 well cell culture plate | Corning | Cat#3526 |
| Sterile 100mm cell culture dish | Corning | Cat#430167 |
| Sterile 25G x 1" Needle | BD | Cat#305125 |
| Sterile 10mL syringe | BD | Cat#302995 |
| 96-well EIA/RIA flat bottom plate | Corning | Cat#3369 |
| round-bottom flow test tubes | VWR | Cat#60819-820 |
| 60mm tissue culture dishes | VWR | Cat#82050-546 |
| Scalpel handle | Integra Miltex | Product# 2970018590 |
| 50ml Centrifuge tubes | VWR | Cat#89039-656 |
| 15ml Centrifuge tubes | VWR | Cat#89039-664 |
| 40um cell strainer | VWR | Cat#76327-098 |
| Filter cloth | Fisher scientific | Cat#08-670-190 |
| 1.5 mL Eppendorf tube | USA Scientific | Item#8700-1525 |
| Dissecting scissors | VWR | Cat# 76457-404 |
| Surgical scissors | VWR | Cat# 470001-842 |
| Tweezers | VWR | Cat# 47743-180 |
| Scalpel handle | Integra Miltex | Product# 2970018590 |
| Scalpel blades | Southmedic | Part ID# 73-0410 |
| Electric shaver | Philips | Product#MG3750/60 |

### EXPERIMENTAL MODEL AND STUDY PARTICIPANT DETAILS

#### Mice and obesity-related psoriasis-like skin

The protocol of mouse studies was approved by the Institutional Animal Care and Use Committee (IACUC) at the University of Iowa. Fabp5^f/f^ mice, Fabp5^-/-^ mice, and their wild-type controls were bred and housed in the animal facility at the University of Iowa. LysM-Cre mice of C57BL/6 background were purchased from Jackson Laboratory. Mice were weaned at 3-4 weeks of age and randomly assigned to different groups for special diet feeding, which included a low-fat diet (LFD, 5% fat from soybean), cocoa butter high-fat diet (HFD, 45% fat), olive oil HFD (45% fat), safflower HFD (45% fat), and fish oil HFD (45% fat). Mouse body weight was monitored every three weeks, and body fat deposition was measured via NMR scanning after six months on special diets.

Following six months of dietary treatment, mice were used for psoriasis induction via imiquimod (IMQ) treatment, as previously described^29^. Briefly, hair was removed from the mice’s back skin using 200mg of Nair (a commercial facial removal cream), and the mice were monitored for another 48 hours. The hair-removed mice were then randomly treated with 8 mg of IMQ cream (Perrigo, containing 5% IMQ) for four consecutive days. Disease severity was scored based on erythema, induration, and desquamation. The cumulative score was calculated as the sum of these three parameters. At the endpoint of the IMQ treatment, mice were sacrificed for further immunophenotypic analysis using flow cytometry in various tissues.

### MODEL DETAILS

#### Single-cell suspension preparation

The preparation of single-cell suspension from the skin epidermis and dermis was conducted following our previous work^29^. Briefly, psoriatic skin lesions from mice were removed. The removed skin was placed with the epidermal side down on the lid of the 60mm culture dish containing 6 ml of cold PBS. Subcutaneous fat was scraped off from the skin using a tweezer and the blunt end of a scalpel. The remaining skin tissue was cut into 0.5 x 1 cm pieces and incubated with 6 ml of 0.25% Trypsin-EDTA solution for 1.5 hours at 32 °C with gentle shaking at 20 rpm. After the incubation, the epidermal and dermal layers were separated with the blunt end of a scalpel. Epidermal single cells were obtained by thoroughly dispersing the epidermis tissue with a pipette until all the fragments passed freely through the tip of a 1ml pipette. The cell suspension was then filtered through a 40 µm cell strainer. The dermis tissue was minced and further digested in the tri-enzyme skin-digesting buffer containing 0.2 mg/mL hyaluronidase, 0.02 mg/mL DNase-1, and 0.5mg/mL collagenase IV in RPMI-1640 medium supplemented with 5% fetal bovine serum, for 45 minutes at 37 °C with gentle shaking 20 rpm. Following the digestion, the dermis tissue was transferred into a 50ml centrifuge tube and vortexed for at least 20 seconds. Dermal single cells were then obtained by a two-step filtration process: first through a 105um mesh filter cloth, followed by a 40 µm cell strainer.

For single-cell suspension from draining lymph nodes (dLNs), both lymph nodes from both sides are collected with removing adherent fats and kept in cold PBS of the 60mm tissue culture dish. dLNs are ground in cold PBS to release single cells. Cell suspension was filtered through a 40µm cell stainer into a 15mL centrifuge tube. Cells were then washed twice with cold PBS, pelleted, and resuspend in cold PBS.

For splenic single-cells, mouse spleen was collected and ground to release single cells. Cells were filtered through a 40µm cell stainer into a 15mL centrifuge tube, and pelleted. Cells were then processed red blood cell lysis by resuspended in 3mL RBC lysis buffer (Tonbo Biosciences) and incubated for 3 minutes at room temperature. Cells were then washed by cold PBS, pelleted, and resuspended in cold PBS.

For peripheral blood single cells, 100uL blood from mice were added into 5mL RBC lysis buffer (Tonbo Biosciences) and incubated at room temperature for 10 minutes. Cells were washed with cold PBS, pelleted, and process 10 minutes of RBC lysis once more. Cells were then washed again, pelleted, and resuspended in cold PBS.

For bone marrow cells, a mouse femur was dissected. Muscle was removed from the femurs, and both ends were cut to flush the bone marrow with cold PBS into a 15mL conical tube using a 10mL syringe with a 25G needle. The Bone marrow cells were washed, filtered through a 40μm cell strainer, resuspended in 1mL red blood cell lysis buffer (Tonbo Biosciences), and incubated at room temperature for 3 minutes. Cells were washed again with cold PBS, pelleted, and resuspended.

#### Flow cytometry

Single-cell suspensions were prepared from various mouse tissues, including the epidermis, and dermis, draining lymph nodes, spleens, blood, and bone marrow. For surface staining, cells were incubated in cold PBS for 30 min with the following flow antibodies: Zombie-violet( for live/death cells), anti-mouse CD45-BUV563, CD11b-BUV737, TCR γ/δ-APC, CD4-BUV395, CD8α-BV615, B220-AF700, F4/80-AF488, CD11c-BV711, NK1.1-PE-Cy5, MHCII-BV605, Ly6C-BB700, Ly6G-Spark blue-550, CD27-BV650, CD3 ε-BV785, CD207-PE, and CD16/32 (Fc blocker). For intracellular cytokine staining, cells were treated with a Cell Activation Cocktail (Biolegend in RPMI-1640 medium containing 5% fetal bovine serum for 4 hours at 37 °C, followed by the surface staining using the following antibodies: CD45-BUV563, CD11b-BUV737, CD4-BUV395, CD8α-BV615, F4/80-AF488, CD11c-BV711, TCR γ/δ -PE-Cy5, CD3 ε-BV785, CD207-PE, and CD16/32 (Fc blocker). Afterward, cells were washed with PBS, fixed, and permeabilized using the intracellular staining buffer set (Cat. #424401, Biolegend) according to the manufacturer’s instructions. Intracellular staining was performed using the following antibodies: anti-mouse FOXP3-AF700, TNFα-APC-Cy7, IL-23-AF647, IFNγ-BV510, IL-17A-BV650, IL-1β-PE-Cy7, IL-4-BV605, and IL-10-Pacific Blue. For intracellular FABP5 staining, cells stained by surface markers underwent the intracellular staining process using anti-mouse FABP5 antibody for 1.5 hours at 4 °C, followed by secondary staining of Donkey-anti-goat IgG-AF594 antibody. Cells were acquired by Cytek Aurora Flow Cytometer.

#### UMAP and FlowSOM analysis of flow cytometry data

Uniform manifold approximation and projection (UMAP) analysis of flow cytometric data was performed using FlowJo v.10.8.1 software (FlowJoLLC). An equal number of mice from each genotype and experimental group were included in the analysis. For each mouse, cells from both the epidermis and dermis were analyzed. Equal cell numbers of CD45^+^Zombie-violet^-^ live immune cells were selected using the DownSample plugin in FlowJo. The selected cells were then merged using the concatenate tool and barcoded to enable further tracking and distinction between samples. UMAP was performed using the markers indicated in the figures, with expression levels represented by color code normalized to the median fluorescence intensity. The levels were displayed on a four-color scale ranging from blue (low), green (intermediate), yellow, to red (high). FlowSOM clustering analysis was conducted on both combined samples from different groups and individual samples from each group. The identified clusters were compared between groups and confirmed through manual gating.

#### Analysis of bulk RNA sequencing and single-cell transcriptomic data

RNA sequencing dataset (GSE161296) from our previous study^4^ was downloaded from the Gene Expression Omnibus (GEO) database. Relative expression levels of individual genes between wild-type and Fabp5^-/-^ mouse skins were extracted from the dataset and analyzed using GraphPad Prism 10 software. Differential gene expression (DEG) analysis was performed as previously described^4^ and visualized with a volcano plot using MaGIC Volcano Plot Tool (https://volcano.bioinformagic.tools/). DEGs were selected based on an adjusted p-value <0.05 and log2 fold change <-0.5. Pathway analysis was conducted using g: Profiler^52^ to identify enriched pathways in the KEGG and Gene Ontology (GO: Biological Process) database. Single-cell transcriptomic datasets included Tabula Sapiens – Skin and blood. The analyses were performed using BxGenomics scRNA-Seq View (https://app.bxgenomics.com/bxg/app/).

#### Cell culture

Fabp5^-/-^ Raw264.7 cells and their wild-type (WT) controls were generated using CRSPR-Cas9 system by Ubigene company (www.ubigene.us). Briefly, a guide RNA (gRNA), G13 (sequence: CCATCCCACAGGAGTAGGACTGG) was designed to target the upstream region of Exon-2 of the Fabp5 gene, creating a double-strand break by the Cas9 enzyme. WT and Fabp5^-/-^ Raw264.7 cells were cultured in RPMI-1640 containing 10% FBS and antibiotics (Cat. #30–2300, ATCC).

To obtain Granulocyte/macrophage-colony stimulating factor (GM-CSF)-driven bone marrow-derived macrophages (GM-BMMs), femurs were dissected from CO_2_-euthanized, age/sex-matched WT and Fabp5^-/-^ mice. Bone marrow cells were obtained as mentioned above and resuspended in 15mL RPMI-1640 with 10% FBS and supplemented with 50ng/mL recombinant mouse GM-CSF (Biolegend), then seeded on 100mm culture dishes with a cell concentration of 5x10^6^/mL. Cells were cultured for 7 days, with half of the medium replaced with fresh GM-CSF-containing medium on day 2 and day 5. On day 7, cells were harvested by scraping and re-seeded in culture plates for further treatment.

For peritoneal macrophages, 6mL of PBS was injected into the peritoneal cavity of CO_2_-euthanized, age/sex-matched WT and Fabp5^-/-^ mice and left to sit for 3 minutes. The skin was carefully cut open without damaging the peritoneum, and PBS from the peritoneal cavity was aspirated using a 10mL syringe with a 25G needle, then transferred to a 15mL conical tube. Cells were pelleted, resuspended in RPMI-1640 with 10% FBS, and counted for in vitro experiments. Cells were seeded into cell culture plates and incubated overnight. Non-adherent cells were washed away with PBS, leaving adherent peritoneal macrophages for further treatment.

#### Free Fatty acid (FFA) preparation and treatment

Due to their low solubility, free fatty acids (FFAs) were conjugated with bovine serum albumin (BSA) as described ^42^. FFAs, including stearic acid (SA), oleic acid (OA), and docosapentaenoic acid (DPA), were purchased from Nu-Chek Prep (MN). Each FFA was conjugated with a pre-prepared solution of 2 mM endotoxin-free, fatty acid-free BSA in PBS at a concentration of 5 mM. The FFA-BSA conjugates were sonicated until fully dissolved, then filtered through a 0.2 μm sterile filter for use in cell culture studies. Conjugates were subsequently added to the cell culture system at the specified final concentration of 200μM.

#### Extracellular ATP Assay

Supernatant from Raw264.7 cells subjected to different treatments was collected. Extracellular ATP levels in the supernatant were measured using a bioluminescence-based assay with the ATP Determination Kit (Thermo Fisher). Briefly, standard samples and experimental supernatant were diluted 1:10 in a reaction solution containing 1 mM DTT, 0.5 mM D-luciferin, and 1.25 ug/ml firefly luciferase. The mixture was gently mixed and incubated for 15 min at 28 °C. Following incubation, luminescence was read by a luminometer in a BioTek Synergy LX Multimode Reader. An ATP standard curve was generated using standard samples, and the ATP levels in the experimental samples were calculated based on this curve.

#### Seahorse Assay

The mitochondrial oxygen consumption rate (OCR) of macrophages was measured using the Seahorse XFe96 Bioanalyzer (Agilent). Macrophages were seeded at a concentration of 5×10⁴ cells per well in RPMI-1640 medium with 10% FBS. Following incubation, cells were washed with RPMI-XF medium without FBS and then maintained in RPMI-XF medium at 37°C. OCR measurements were taken at basal levels and following sequential additions of 200 µM BSA or fatty acids, 2.5 µM oligomycin, 1 µM FCCP, and 1 µM Rotenone/Antimycin A. Assays were conducted on the XFe96 extracellular flux analyzer, with automated normalization based on live cell counts at the conclusion of the assay.

#### Quantitative real-time PCR

Supernatants from cultured macrophages were discarded, and cells were washed twice with cold PBS and lysed using the Lysis Buffer from the PureLink RNeasy Mini Kit (Cat. #12183025, Invitrogen, Waltham, MA). RNA was then purified following the manufacture’s instructions. cDNA synthesis was performed using a QuantiTect Reverse Transcription Kit (Cat. #205314, Qiagen, Hilden, Germany). Real-time PCR was conducted with Power SYBR Greed PCR Master Mix (Cat. #4368708, Applied biosystems, Waltham, MA) on a StepOnePlus qPCR Systems (Applied Biosystems). Primers for target genes, including Fabp5, IL-1β, inflammasome-related genes, FAO-related genes, and housekeeping genes, were obtained from IDT (see Table S1). Relative mRNA levels were calculated by ΔΔCt method using hprt1 as the internal control.

#### PAGE and Western blot

Supernatants from cultured GM-BMMs were discarded, and cells were washed twice with cold PBS followed by lysis with RIPA Buffer (Cat. #9806S, Cell Signaling Technology). Proteins were extracted from the lysate by centrifugation at maximum speed, followed by supernatant collection. Protein concentrations were measured in each sample using BCA assay (Cat. #23225, Thermo Scientific). The samples were then mixed with SDS loading buffer containing 2-mercaptoethanol, boiled at 90°C for 5 min, and loaded to an SDS-PAGE gel to be separated. After SDS-PAGE, proteins were transferred to polyvinylidene difluoride membranes. Target proteins were probed with specific antibodies, including anti-mouse Caspase-1 (CST, Cat. #24232) and anti-mouse cleaved-caspase-1 (CST, Cat. #89332) with β-actin (Thermo Fisher Cat. # MA1-140-A488) as the loading control. A Bio-Rad imaging system was used to visualize and quantify protein levels.

#### ELISA

IL-1β secreted by cultured macrophages were analyzed using the ELISA MAX™ Deluxe Set Mouse IL-1β (Cat. #432604, Biolegend) according to the manufacturer’s instructions. Briefly, capture antibodies for IL-1β was pre-coated in a 96-well flat-bottom ELISA plate (Corning, Cat. #3369). A total of 100μL of each standard and supernatant from the cultured cells were added to the wells, incubated with the capture antibodies, washed, incubated with detection antibodies, washed again, followed by incubation with Avidin-HRP reagent, additional washes, and development by TMB substrate. The plate was read at 450nm and 550nm a spectrophotometer. A standard curve was generated using the standard wells, and the exact IL-1β concentrations were calculated based on this curve.

#### Quantification and statistical analysis

All data were presented as the mean ± SD. All experiments were performed by at least three independent replicates. For both in vitro and in vivo experiments, two-tailed, unpaired Student’s t-test, two-tailed unpaired multiple t-test, and one-way ANOVA followed by Bonferroni’s multiple comparisons test were used. Statistical significance was defined as ∗p < 0.05, ∗∗p < 0.01, ∗∗∗p < 0.001, and ∗∗∗∗p < 0.0001, ns, non-significant.

## Supplemental Figure S1–S7 and Legends

**Figure S1.**
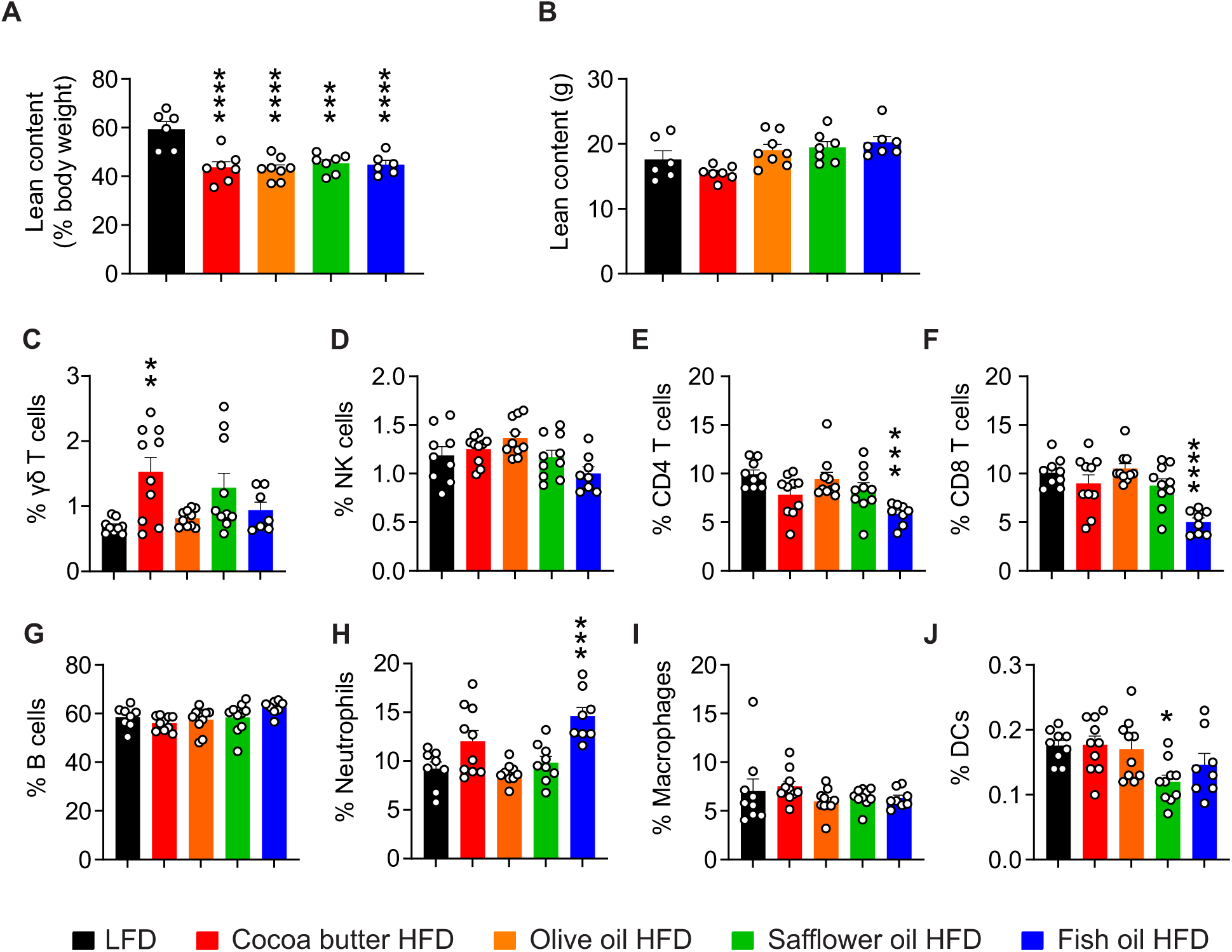
HFDs of different sources altered the systematic immune system, related to Figure 1. A-B. NMR scanning for lean body content in mice fed customized HFDs and LFD for 6 months, shown as a ratio (A) and in weight (B). Statistical analysis of the averages was performed on each HFD groups compared to LFD group. C-J. Flow cytometric analysis of peripheral blood immunophenotypes of mice fed customized HFDs and LFD for 6 months. Statistical analysis of the averages was performed on each HFD group compared to LFD group. Data are pooled from two independent experiments with 4-5 mice per group in each experiment (A-J). Data are shown as mean ± SEM ∗p ≤ 0.05 ∗∗p ≤ 0.01, ∗∗∗p ≤ 0.001, ∗∗∗∗p ≤ 0.0001, ns, non-significant, one-way ANOVA with Bonferroni’s multiple comparisons test.

**Figure S2.**
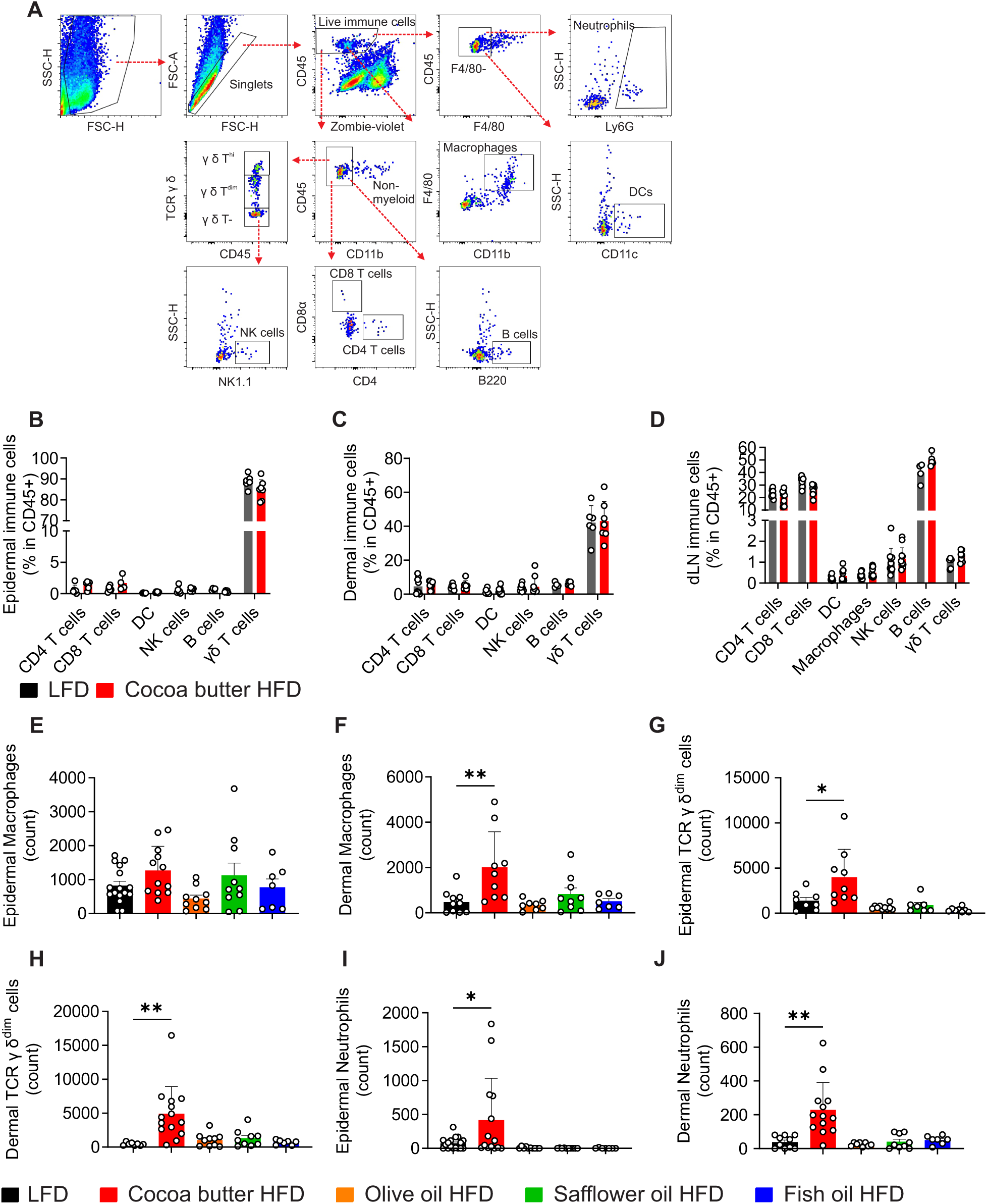
Saturated, not unsaturated HFD promotes proinflammatory immune cell infiltration in the skin of IMQ-psoriasis, related to Figure 2. A. Proposed gating strategy for immunophenotypic analysis of psoriatic skin tissue from mice. B-D. Immunophenotypic staining of epidermis (B), dermis (C), and draining lymph nodes (D) from mice fed cocoa butter HFD or LFD, treated with IMQ for 4 days. E-J. Flow cytometric analysis of count of epidermal and dermal macrophages (E, F), TCR γ δdim cells (G, H), and neutrophils (I, J) on custom-made HFDs, treated with IMQ for 4 days. Data are pooled from two independent experiments with 3-5 mice per group (B-J). Data are shown as mean ± SEM ∗p ≤ 0.05 ∗∗p ≤ 0.01, ∗∗∗p ≤ 0.001, ∗∗∗∗p ≤ 0.0001, ns, non-significant, unpaired multiple t-test for panels B-D, or one-way ANOVA with Bonferroni’s multiple comparisons test for panels E-J.

**Figure S3.**
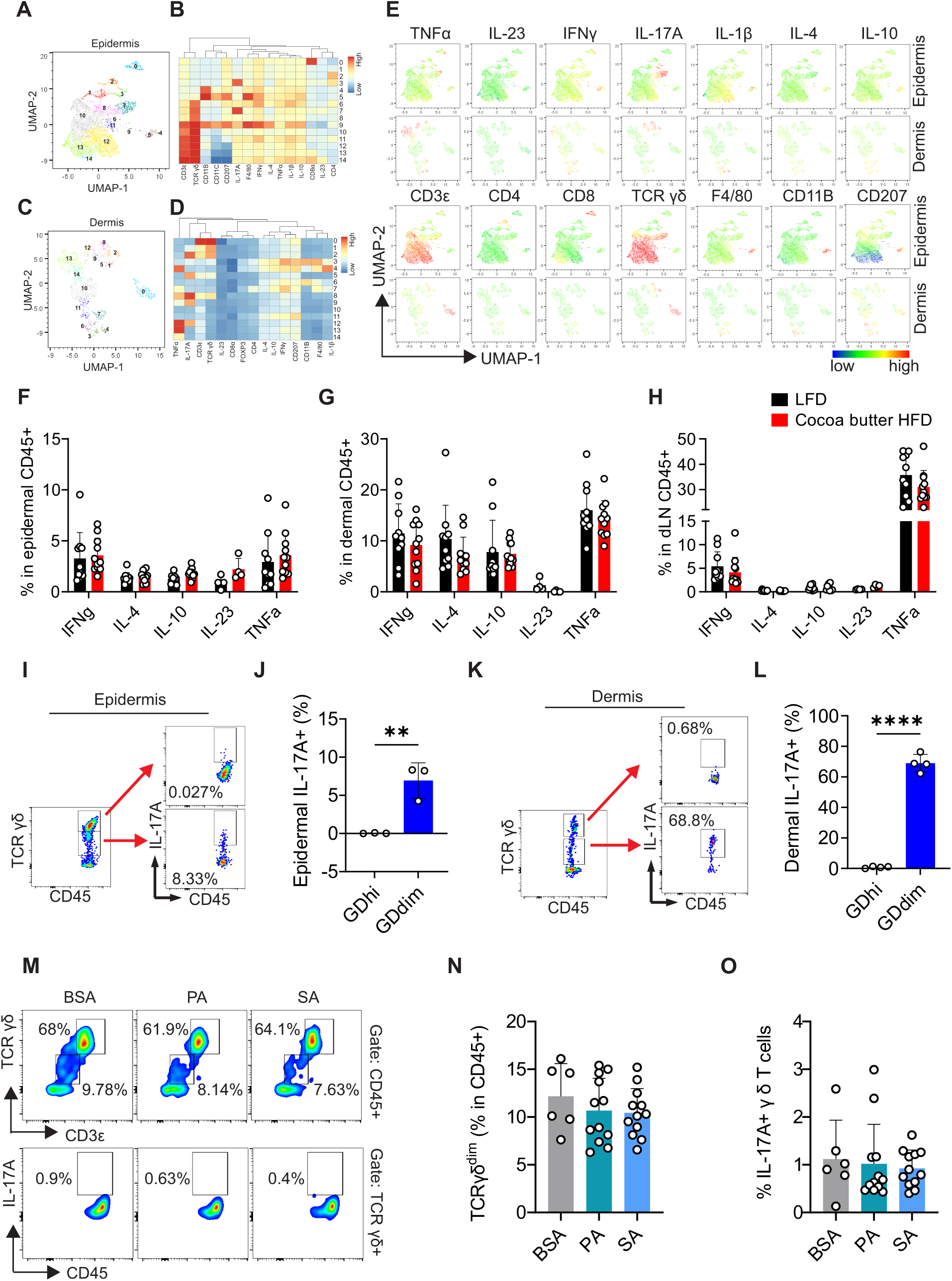
Saturated HFD amplifies skin IL-1β/IL-17A signaling in psoriasis, related to Figure 3. A-E. Representative surface and intracellular flow staining of immune cells from epidermis and dermis after 4 days of IMQ treatment. UMAP visualization (A, C) and heatmap (B, D) of 15 immune cell clusters analyzed by FlowSOM. (E) Individual marker signatures. F-H. Flow cytometric analysis of cytokine expression in immune cells from the epidermis (F), dermis (G), and dLNs (H) of mice fed cocoa butter HFD or LFD, treated with IMQ for 4 days. I-L. IL-17A expression levels of TCR γ δ dim cells and TCR γ δ hi cells from the epidermis (I, J) and dermis (K, L) of normal C57BL/6 mice analyzed by flow cytometry. M-O. Epidermal and dermal single-cell suspension mixture from normal mouse skin was treated in vitro overnight with PA, SA, or BSA. Ratios of TCR γ δ dim cells in CD45+ cells and IL-17A+ in γ δ T cells (M) were analyzed by flow cytometry. Statistical analysis was shown in (N) and (O). Combination of both dermis and epidermis from n=3-4 mice from both LFD and cocoa butter HFD groups was included in panels A-E. Data are pooled from two independent experiments for panels F-H, and N-O. Data are shown as mean ± SEM, ∗∗p ≤ 0.01, ∗∗∗∗p ≤ 0.0001, unpaired two-tailed multiple t-test for panels F-H, unpaired Student’s t-test for panels J and L, or one-way ANOVA with Bonferroni’s multiple comparisons test for panels N-O.

**Figure S4.**
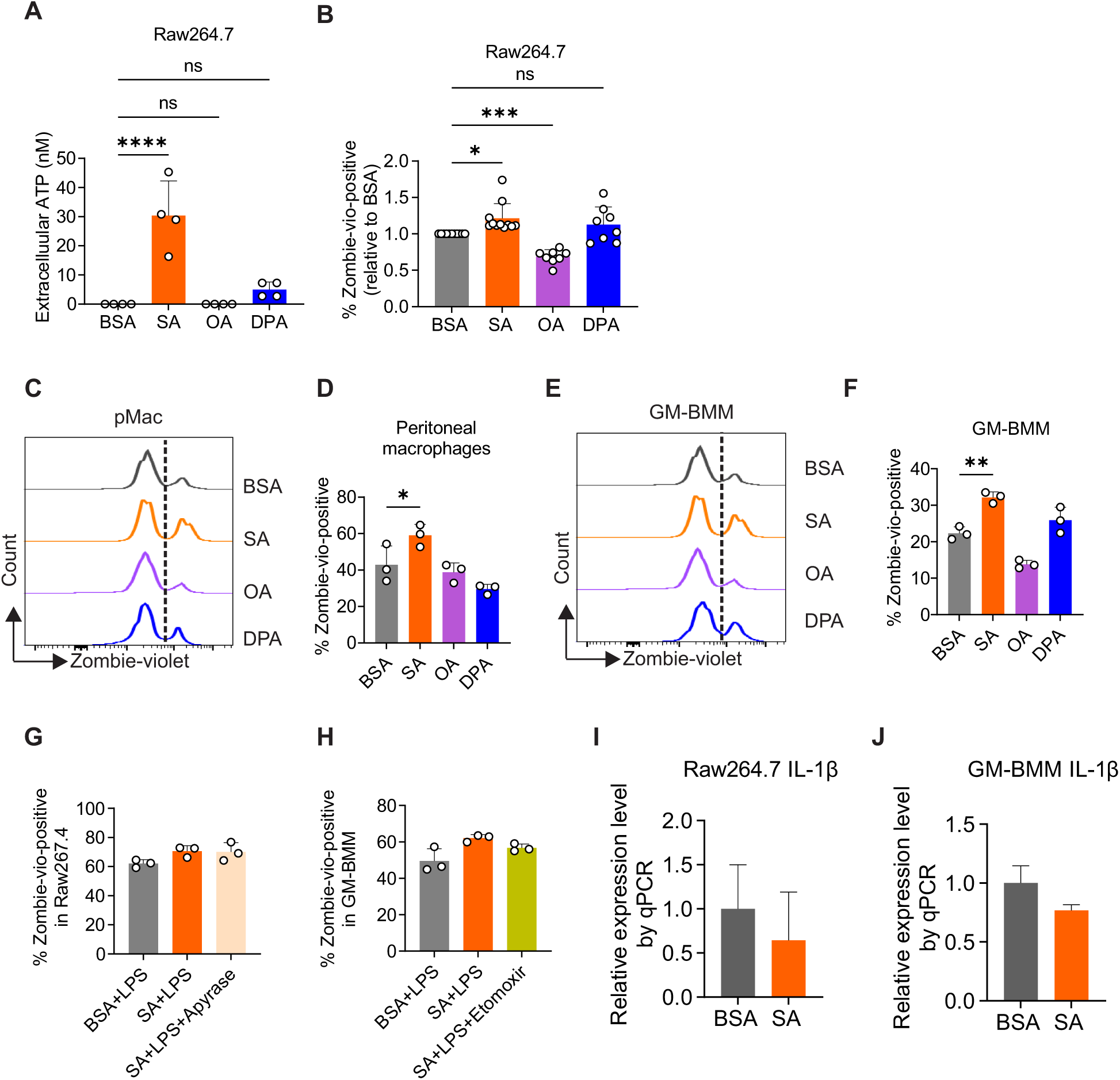
Saturated, not unsaturated FA induces macrophage cell membrane disruption and ATP release, related to Figure 4. A. Extracellular ATP concentration in the supernatant of Raw264.7 cells treated with FAs or BSA in the absence of LPS for 6 hours. B. Raw264.7 cells were treated with FAs or BSA in the absence of LPS for 12 hours. Cell membrane disruption was assessed by Zombie-violet staining and analyzed by flow cytometry. C-F. Peritoneal macrophages (C, D) and GM-BMMs (E, F) were treated with SA or BSA in the presence of LPS for 12 hours. Cell membrane disruption was assessed by Zombie-violet staining and analyzed by flow cytometry. G. Cell membrane disruption of Raw264.7 cells treated with SA or BSA in the presence of LPS for 12 hours, with or without 5 units/ml Apyrase, and analyzed by Zombie-violet staining. H. Cell membrane disruption of GM-BMMs treated with SA or BSA in the presence of LPS for 12 hours, with or without 200uM Etomoxir, analyzed by Zombie-violet staining. I-J. Expression levels of IL-1β genes in Raw264.7 cells (I) and GM-BMMs (J) treated by SA or BSA for 4 hours, analyzed by real-time PCR. Data are shown as mean ± SEM, ∗p ≤ 0.05, ∗∗p ≤ 0.01, ∗∗∗p ≤ 0.001, ∗∗∗∗p ≤ 0.0001, ns, non-significant, one-way ANOVA with Bonferroni’s multiple comparisons test for panels A, B, D, F, G, and H, or unpaired two-tailed Student’s t-test for panels I-J.

**Figure S5.**
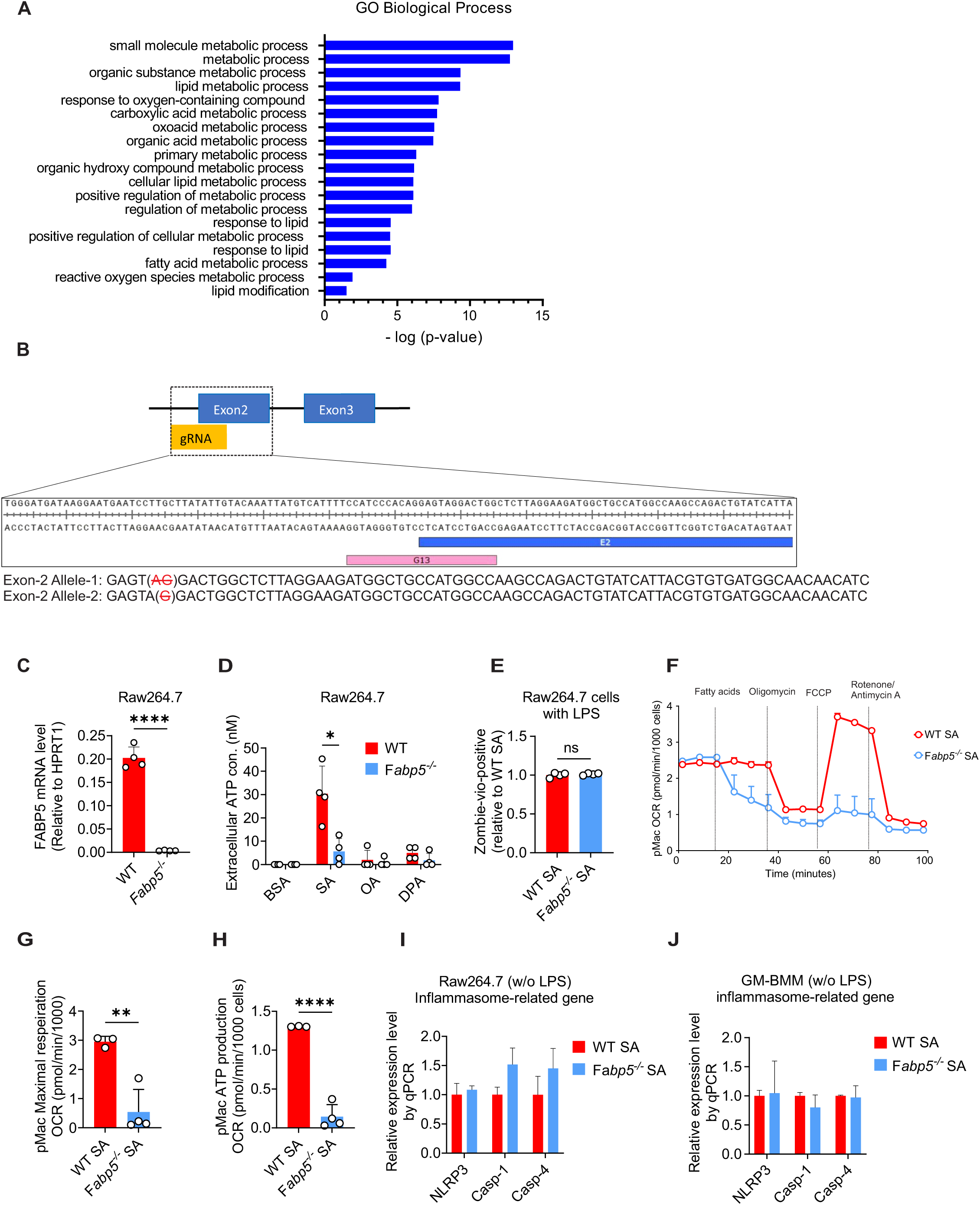
FABP5 is critical for macrophage FAO and ATP production by saturated FA, related to Figure 5. A. Analysis of gene oncology biological process from DEGs with an adjusted p value cutoff of 0.05 and a Log2 fold change cutoff of -0.5. B. Schematic of the guide-RNA (gRNA) design Fabp5-/-Raw264.7 cells generation using CASPR-Cas9 system. C. Real-time PCR analysis of FABP5 mRNA levels of both WT and Fabp5-/-Raw264.7 cells. D. Supernatant ATP concentration of WT and Fabp5-/-Raw264.7 cells treated with FAs or BSA in the absence of LPS for 6 hours. E. Cell death of WT and Fabp5-/-Raw264.7 cells treated with SA in the presence of LPS for 12 hours analyzed by Zombie-violet staining. F-H. OCR of peritoneal macrophages isolated from WT and Fabp5-/-mice was measured using Seahorse (F), with statistical analysis of maximal respiration OCR (G) and ATP production (H). I-J. Inflammasome-related gene expression of WT and Fabp5-/-Raw264.7 cells (I) and GM-BMMs (J) treated with 200µM BSA or SA for 4 hours, analyzed by real-time PCR. Data are shown as mean ± SEM, ∗p ≤ 0.05, ∗∗p ≤ 0.01, ∗∗∗∗p ≤ 0.0001, ns, non-significant, unpaired two-tailed Student’s t-test for panels C, E, G, and H, or unpaired two-tailed multiple t-test for panels D, I, and J.

**Figure S6.**
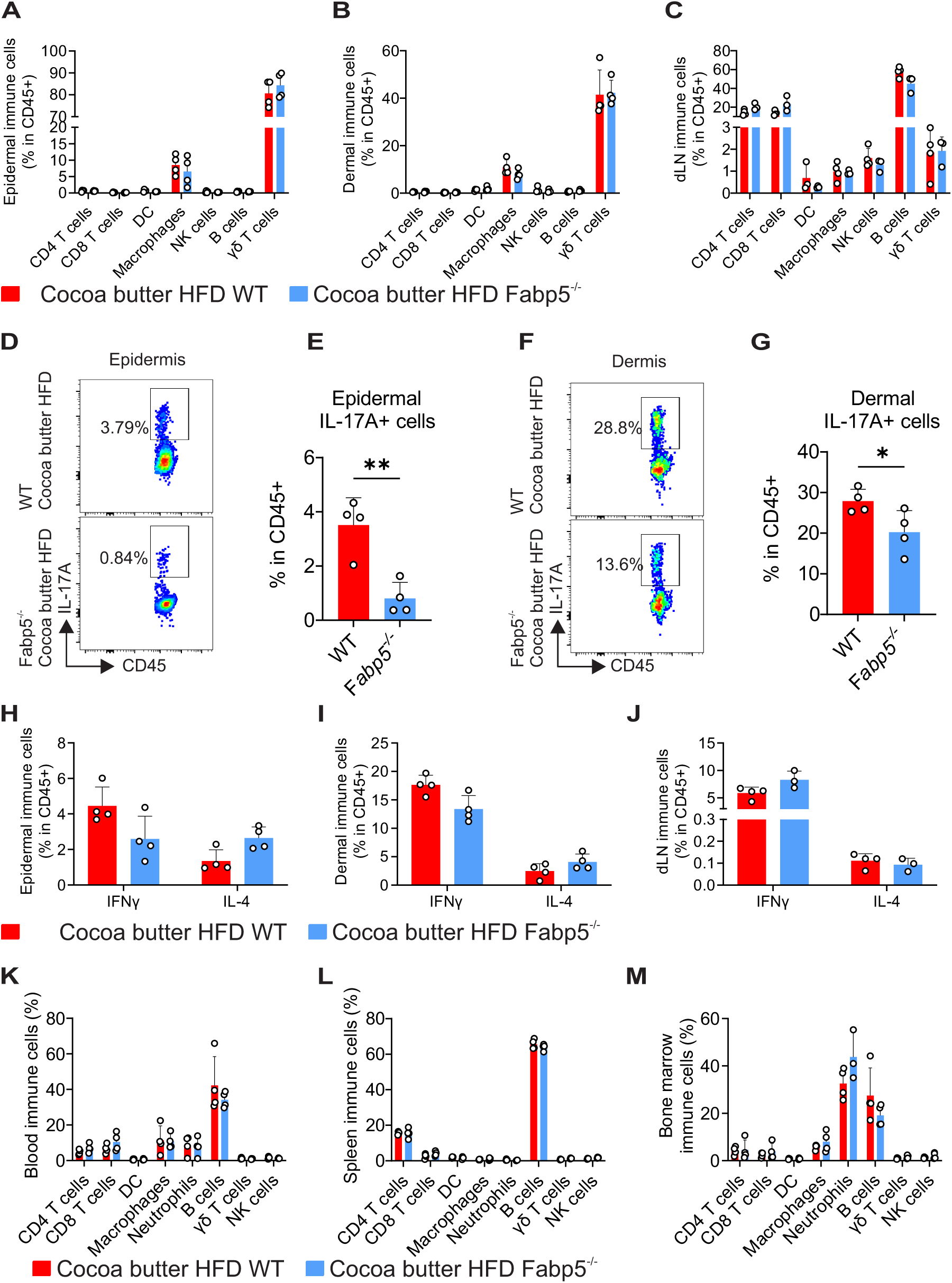
Immunophenotypic flow staining of Fabp5^-/-^ mice compared to WT mice in saturated HFD-related psoriasis, related to Figure 6. A-C. Flow cytometric analysis of immunophenotypes of the epidermis (A), dermis (B), and dLNs (C) in IMQ-induced psoriasis of WT and Fabp5-/-mice fed cocoa butter HFD. D-G. Flow cytometric analysis of ratio of epidermal (D-E) and dermal (F-G) IL-17A+ immune cells of WT and Fabp5-/-mice fed cocoa butter HFD, treated with IMQ for 4 days. H-J. Flow cytometric analysis of cytokine expression in CD45+ immune cells in the epidermis (H), dermis (I), and dLNs (J) in IMQ-induced psoriasis of WT and Fabp5-/-mice fed cocoa butter HFD or soybean LFD. K-M. Flow cytometric analysis of immunophenotypes in peripheral blood (K), spleen (L), and bone marrow (M) in IMQ-induced psoriasis of WT and Fabp5-/-mice fed cocoa butter HFD or soybean LFD. Data are shown as mean ± SEM, ∗p ≤ 0.05, ∗∗p ≤ 0.01, unpaired two-tailed multiple t-test for panels A-C, H-J, and K-M, or unpaired two-tailed Student’s t-test for panels E and G.

**Figure S7.**
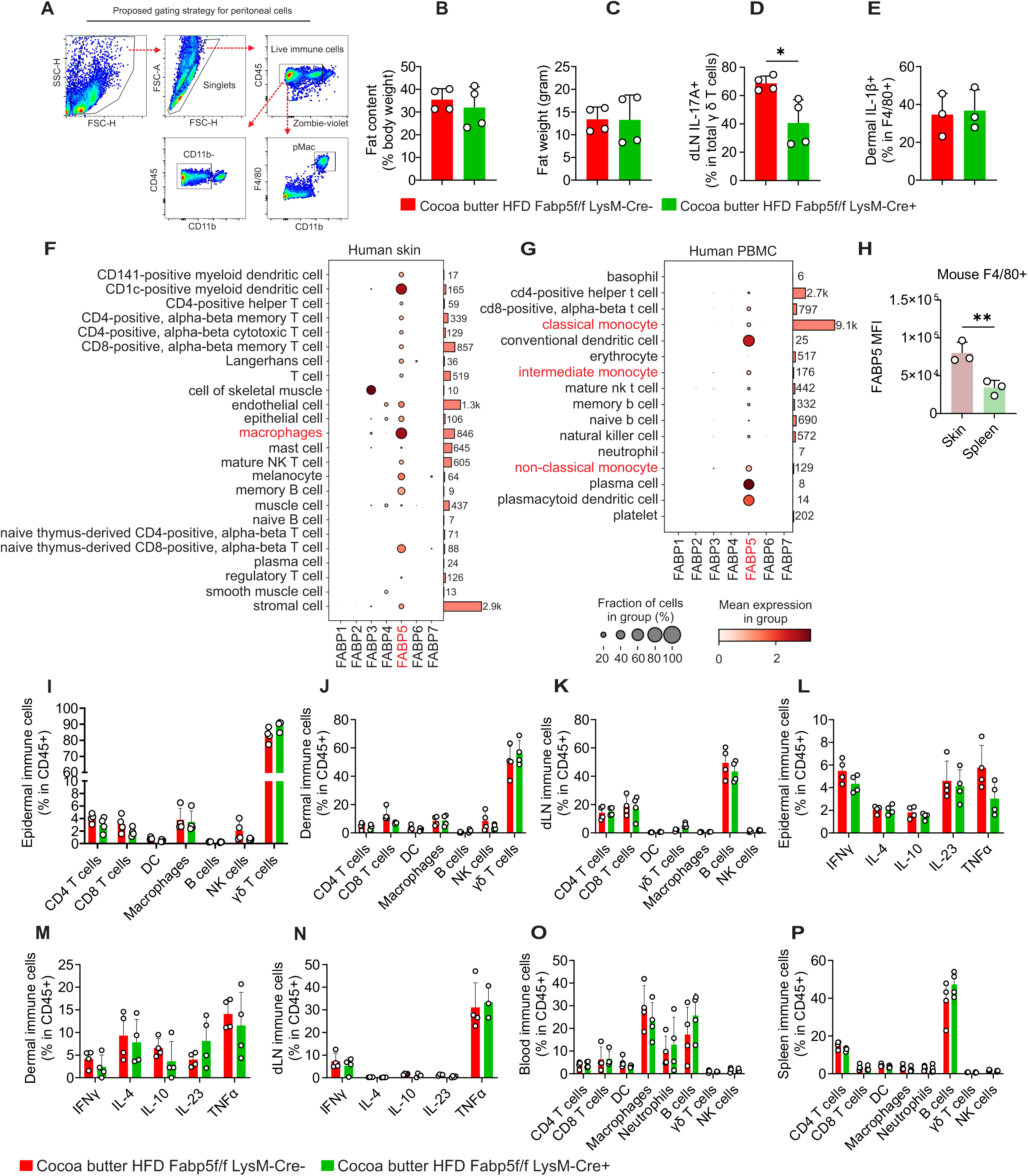
immunophenotypic flow staining of macrophage-specific FABP5-deficient mice compared to WT mice in saturated HFD-related psoriasis, related to Figure 7. A. Proposed gating strategy for peritoneal macrophages and CD11b-peritoneal immune cells. B-C. NMR scanning for body fat content of Fabp5f/f LysM-Cre-and Fabp5f/f LysM-Cre+ mice fed cocoa butter HFD or LFD for 6 months. D. Flow cytometric analysis of ratio of IL-17A+ among γ δ T cells in dLNs of Fabp5f/f LysM-Cre- and Fabp5f/f LysM-Cre+ mice fed cocoa butter HFD, treated with IMQ for 4 days. E. Flow cytometric analysis of IL-1β+ in F4/80+ macrophages in the dermis of Fabp5f/f LysM-Cre- and Fabp5f/f LysM-Cre+ mice on cocoa butter HFD, treated with IMQ for 4 days. F-G. Analysis of the expression profile of FABP family members in different immune cells from human skin (F) and PBMC (G) using a publicly accessible dataset GSE201333. H. Flow cytometric analysis of FABP5 expression level in mouse F4/80+ cells from skin and spleen. I-K. Immunophenotypic flow staining of the epidermis (I), dermis (J), and dLNs (K) from Fabp5f/f LysM-Cre- and Fabp5f/f LysM-Cre+ mice on cocoa butter HFD, treated with IMQ for 4 days. L-N. Intracellular flow staining for cytokine expression in CD45+ cells from the epidermis (L), dermis (M), and dLNs (N) of Fabp5f/f LysM-Cre- and Fabp5f/f LysM-Cre+ mice on cocoa butter HFD, treated with IMQ for 4 days. O-P. Immunophenotypic flow staining of peripheral blood (O) and spleen (P) from Fabp5f/f LysM-Cre- and Fabp5f/f LysM-Cre+ mice on cocoa butter HFD, treated with IMQ for 4 days. Data are shown as mean ± SEM, ∗p ≤ 0.05, ∗∗p ≤ 0.01, unpaired two-tailed Student’s t-test for panels B-E and H, or unpaired two-tailed multiple t-test for panels I-P.

## Supplemental Table S1

**Supplemental Table S1.**
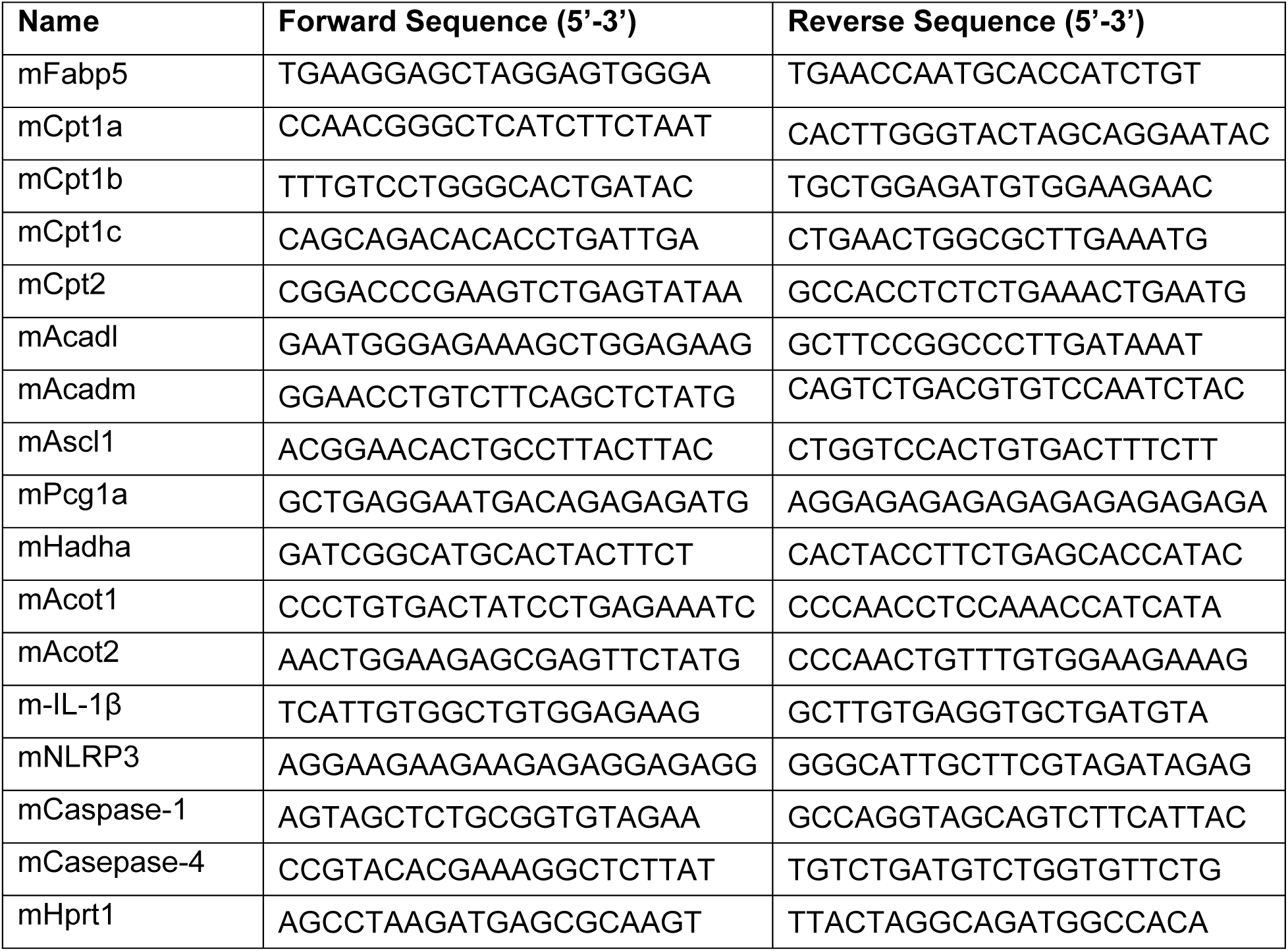
Real-time PCR primers.

